# Epigenetic repression of Wnt receptors in AD: a role for Sirtuin2-induced H4K16ac deacetylation of Frizzled1 and Frizzled7 promoters

**DOI:** 10.1101/2021.05.19.444683

**Authors:** Ernest Palomer, Núria Martin-Flores, Sarah Jolly, Patricia Pascual-Vargas, Stefano Benvegnù, Marina Podpolny, Samuel Teo, Kadi Vaher, Takashi Saito, Takaomi C Saido, Paul Whiting, Patricia C Salinas

## Abstract

Growing evidence supports a role for deficient Wnt signalling in Alzheimer′s disease (AD). First, the Wnt antagonist DKK1 is elevated in AD brains and is required for amyloid-β-induced synapse loss. Second, LRP6 Wnt co-receptor is required for synapse integrity and three variants of this receptor are linked to late-onset AD. However, the expression/role of other Wnt signalling components remain poorly explored in AD. Wnt receptors Frizzled1 (Fzd1), Fzd5, Fzd7 and Fzd9 are of interest due to their role in synapse formation/plasticity. Our analyses showed reduced *FZD1* and *FZD7* mRNA levels in the hippocampus of human early AD stages and in the hAPP^NLGF/NLGF^ mouse model. This transcriptional downregulation was accompanied by reduced levels of the pro-transcriptional histone mark H4K16ac and a concomitant increase of its deacetylase Sirt2 at *Fzd1* and *Fzd7* promoters in AD. *In vitro* and *in vivo* inhibition of Sirt2 rescued *Fzd1* and *Fzd7* mRNA expression and H4K16ac levels at their promoters. In addition, we showed that Sirt2 recruitment to *Fzd1* and *Fzd7* promoters is dependent on FoxO1 activity in AD, thus acting as a co-repressor. Finally, we found reduced levels of Sirt2 inhibitory phosphorylation in nuclear samples from human early AD stages with a concomitant increased in the Sirt2 phosphatase PP2C. This results in hyperactive nuclear Sirt2 and favours *Fzd1* and *Fzd7* repression in AD. Collectively, our findings define a novel role for nuclear hyperactivated Sirt2 in repressing *Fzd1* and *Fzd7* expression *via* H4K16ac deacetylation in AD. We propose Sirt2 as an attractive target to ameliorate AD pathology.

## Introduction

Alzheimer’s disease (AD) is the most common form of dementia, clinically characterised by progressive cognitive impairment and memory loss. One of the early events in AD is the loss of synapses, a process strongly correlated with cognitive decline [1, 2]. Interestingly, several signalling pathways required for synapse function and integrity are dysregulated in AD [3, 4]. Of particular interest is the Wnt signalling pathway(s). First, the secreted Wnt antagonist DKK1 is elevated both in the brain of AD patients and models [5–7] and by exposure to Aβ-oligomers (Aβo) [8]. Importantly, blockade or knockdown of Dkk1 protects synapses against Aβ [8, 9]. Second, three genetic variants of the Wnt co-receptor Low-Density Lipoprotein Receptor-Related Protein-6 (LRP6) have been linked to late-onset AD (LOAD) [10, 11]. Third, Wnt3a and Wnt5a are protective against Aβ [12, 13]. Fourth, induced Dkk1 expression in the adult mouse hippocampus leads to synapse loss, plasticity defects and cognitive impairment [14, 15], features that can be reversed by cessation of Dkk1 expression [14]. Together, these findings demonstrate that Aβ deregulates Wnt signalling and that boosting Wnt signalling could be protective to synapses in AD.

Deregulation of Wnt signalling in AD could be mediated by different mechanisms; elevation of Wnt antagonists, such as Dkk1, or down-regulation of key Wnt components such as Wnt proteins or their receptor Frizzled (Fzd) [16]. Interestingly, several Fzd receptors are sufficient and/or required for synaptic assembly. For example, Fzd1 and Fzd5 are involved in the formation of presynaptic terminals whereas Fzd7 and Fz9 promote postsynaptic assembly [17–20] (Fig. 1A). Fzd7 is also required for synaptic plasticity [17]. However, little is known about how these Fzd receptors are regulated in AD.

**Figure 1:**
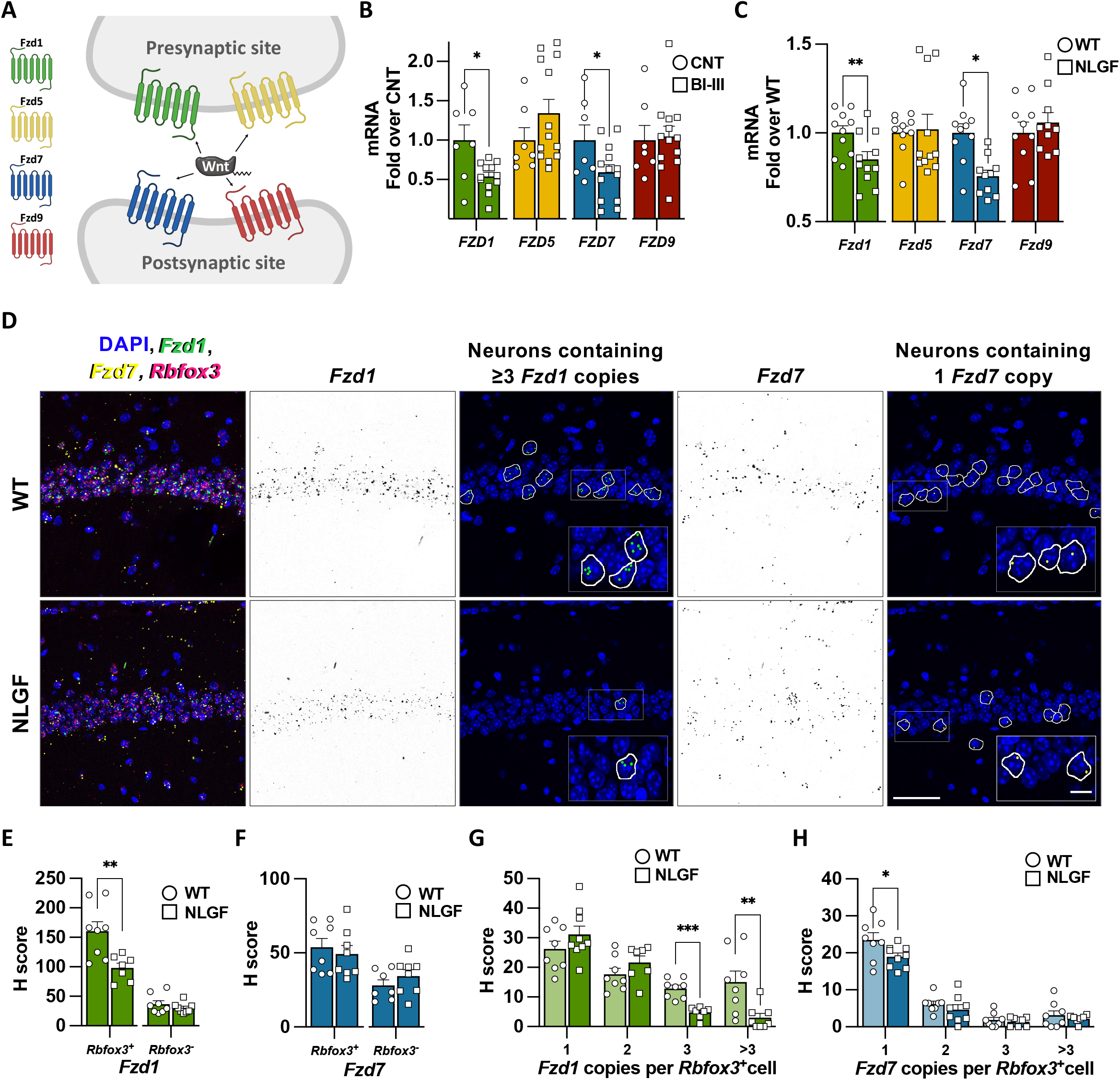
Frizzled 1 and Frizzled 7 are downregulated in early AD. A) Scheme representing a synapse showing the localization at pre- and/or post-synaptic terminal for Fzd1, Fzd5, Fzd7 and Fzd9. B) qPCR analyses showing reduced mRNA levels of *FZD1* and *FZD7* in human hippocampal samples from Braak stages I-III (BI-III) subjects compared to controls. No changes are observed for *FZD5* and *FZD9* in early AD. C) qPCR analysis showing the reduced mRNA levels of *Fzd1* and *Fzd7* in 2-month-old hAPP^NLGF/NLGF^ hippocampus (NLGF). No differences in *Fzd5* and *Fzd9* levels are observed in NLGF hippocampal mRNA. D) Representative smFISH images of WT and NLGF CA1 hippocampal region. First column shows merged images with DAPI (blue), *Fzd1* (green), *Fzd7* (yellow) and *Rbfox3* (magenta) mRNAs. *Fzd1* in black (second column) and its representative neuronal *Rbfox3*^+^ cells corresponding to 3 and >3 *Fzd1* copies (third column). *Fzd7* in black (fourth column) and its representative cells corresponding to one *Fzd7* copy (fifth column). Scale bars represent 50 µm and 12.5 µm in the zoomed in inserts. E-F) Single-cell analyses expressed as H-score for *Fzd1* (E) and *Fzd7* (F) in neuronal (*Rbfox3*^+^) and non-neuronal (*Rbfoz3*^-^) cells. G-H) Single-cell distribution of neurons (*Rbfox3*^+^) containing 1, 2, 3 or >3 transcripts for *Fzd1* (G) or *Fzd7* (H). Data are represented as mean + SEM. Statistical analysis by *t-*Test in B for *FZD1, FZD7* and *FZD9* and by Mann-Whitney for *FZD5*; in C *t-*Test for all genes analysed; in E and F *t-*Test for neuronal and non-neuronal; in G *t-*Test for 1, 2 and 3 copies and by Mann-Whitney for >3 copies; in H *t-*Test for 1 and 2 copies and by Mann-Whitney for 3 and >3 copies. N are indicated in each bar by the number of symbols. Asterisks indicate \**p*<0.05; \*\**p*<0.01, \*\*\**p*<0.005.

Here, we investigated whether Fzd receptors are deregulated in AD and the mechanisms involved. We found that *Fzd1* and *Fzd7* are downregulated in early AD by a shared epigenetic mechanism depending on nuclear Sirtuin2 (Sirt2) hyperactivity. We demonstrated that nuclear Sirt2 is recruited to *Fzd1* and *Fzd7* promoters in a FoxO1 dependent manner, leading to reduced levels of the active histone mark H4K16ac at their promoters resulting in their transcriptional repression.

## MATERIAL and METHODS

### Human Tissue

Anonymised human samples were obtained from the Cambridge Brain Bank (CBB) and the Queen Square Brain Bank (QSBB), with informed consent under CBB license (NRES 10/HO308/56) and QSBB licence (NRES 08/H0718/54). Further information can be found in supplementary methods and Table S1.

### Animals

All procedures involving animals were conducted according to the Animals Scientific Procedures Act UK (1986) and in compliance with the ethical standards at University College London (UCL). Further information can be found in supplementary methods.

### Statistical analysis

All values are presented as mean + SEM. Statistical analyses were performed using SPSS v25 (IBM). Outliers were determined with the explore tool (Tukey’s method). Data normality and homogeneity of variances were tested by the Shapiro-Wilk and Levene tests, respectively. Mann-Whitney U-test (two groups) or Kruskal-Wallis followed by Dunn’s multiple comparison (more than two groups) tests were used for non-normally distributed datasets. For normally distributed data; one-sample t-test (two groups with control values equal one), Student’s t-test (two groups) or two-way ANOVA (more than two groups) followed by post-hoc comparisons assuming (Tukey’s) or not assuming equal variances (Games-Howell). All statistical analyses are two-tailed, unless indicated otherwise in the corresponding figure legend. In the figures, asterisks indicate p values as follows: ^*^p < 0.05; ^**^;p < 0.01; ^***^p < 0.005.

## RESULTS

### *Frizzled1* and *Frizzled7* expression is downregulated in AD

Deficient Wnt signalling has been linked to AD by studies on the Wnt antagonist DKK1 and LRP6 genetic variants [21]. In addition, Wnt ligands have been shown to be protective against Aβ insult [21]. However, very little is known about the regulation of Frizzled receptors (Fzd) in AD. Four Fzd receptors (Fzd1, Fzd5, Fzd7 and Fzd9) have been shown to regulate synapse formation and/or function [17–20] (Fig. 1A). We therefore evaluated the expression levels of these receptors. We performed RT-qPCR on human hippocampal RNA samples from control and from subjects with early Braak stages but no cognitive deficits (BI-III; Table S1). We found reduced *FZD1* and *FZD7* mRNA levels in BI-III samples (Fig. 1B). In contrast, *FZD5* and *FZD9* mRNA levels were unchanged (Fig. 1B). These results suggest that two *FZDs* with synaptic function are downregulated in early stages of AD.

Next, we investigated whether the mRNA levels of these *Fzds* were also affected in an AD model. We used the knock-in AD line hAPP^NLGF/NLGF^ (NLGF), which carries the humanized form of APP with the Swedish, Iberian and Arctic mutations, leading to Aβ overproduction [22]. We analysed *Fzd* expression in hippocampal samples of NLGF animals at 2-months-old, an age when Aβ plaques start to appear [22]. Our results showed reduced levels of *Fzd1* and *Fzd7* expression in NLGF samples, whereas *Fzd5* and *Fzd9* remained unchanged (Fig. 1C). Together, these results demonstrate that *Fzd1* and *Fzd7* expression were reduced in both human BI-III subjects and the AD mouse model at an early disease stage.

Fzds are expressed by different brain cells, including neurons, astrocytes and microglia [23, 24] (Fig. S1A). We therefore asked whether reduced mRNA levels of *Fzd1* and *Fzd7* were neuronal specific. We performed single molecule RNA fluorescent in-situ hybridisation (smFISH) for *Fzd1* and *Fzd7* in the CA1 area of the hippocampus (Fig. 1D). Single-cell analyses revealed reduced *Fzd1* levels in NLGF neuronal cells (Rbfox3^+^), without changes in non-neuronal cells (Rbfox3^-^; Fig. 1E) when compared to control animals. However, no changes in the overall levels of *Fzd7* were observed (Fig. 1F). Next, we analysed the distribution of transcript copy number in neuronal cells. We found that neurons containing ≥3 *Fzd1* transcripts were reduced in AD (Fig. 1G). Interestingly, we observed a reduced number of neurons containing one *Fzd7* transcript in the NLGF (Fig. 1H). The lack of difference in H-score for *Fzd7* could be explained by the lower weighting for percentage of cells with 1 copy (see methods). Together, our results demonstrate that *Fzd1* and *Fzd7* RNA levels are reduced in both human BI-III and AD mouse model, with a clear downregulation of neuronal *Fzd1* expression and reduced number of neurons containing one *Fzd7* transcript in the AD mouse hippocampus.

### *Fzd1* and *Fzd7* promoters present reduced H4K16ac levels with concomitant increase of Sirt2 in AD

The reduced levels of *FZD1* and *FZD7* expression in the human brain at early AD stages led us to hypothesise that a shared epigenetic regulation could contribute to their dysregulation. A previous study showed that the pro-transcriptional histone mark acetylated Histone H4 Lysine 16 (H4K16ac) is enriched at promoters of several Wnt signalling pathway components [25]. Chromatin immunoprecipitation (ChIP)-qPCR showed high levels of H4K16ac, and concomitant low levels of total H4, at actively transcribed genes *Actb* and *Eif5* (Fig S1B-C), which have high levels of H4K16ac in the human brain [26]. In contrast, the repressed genes *Hoxa1* and *Krt16* exhibited low levels of H4K16ac and high levels of H4 (Fig. S1B-C)[26]. Higher levels of H4K16ac were found at *Fzd1* and *Fzd7* promoters than at *Fzd5* and *Fzd9* promoters (Fig S1B-C), suggesting that H4K16ac is enriched at the *Fzd1* and *Fzd7* promoters and might contribute to their regulation.

Next, we analysed H4K16ac levels in human hippocampal samples. First, we found that H4K16ac levels were not altered by the post-mortem interval (PMI) time (Fig. S1D). ChIP-qPCR experiments showed that H4K16ac was reduced at *FZD1* and *FZD7* promoters in BI-III (Fig. 2A-B), whereas no changes were observed at the promoters of our internal controls *FZD5* and *FZD9* (Fig. 2A) or external controls genes *Actb, Eif5, Hoxa1* or *Krt16* (Fig. S1E), collectively referred here as control genes. Reduced H4K16ac levels at *Fzd1* and *Fzd7* promoters were also observed in the NLGF hippocampus (Fig. 2C, S1F). Changes in H4K16ac levels could arise from nucleosome remodelling or from differential levels of H4K16ac *per se*. We found no changes in nucleosome remodelling when analysed by H4 total levels, thus changes of H4K16ac at *FZD1* and *FZD7* promoters were likely due to reduced H4K16ac levels (Fig. S1G-H). Together, these results show reduced H4K16ac levels, which could contribute to *FZD1* and *FZD7* repression in early AD.

**Figure 2:**
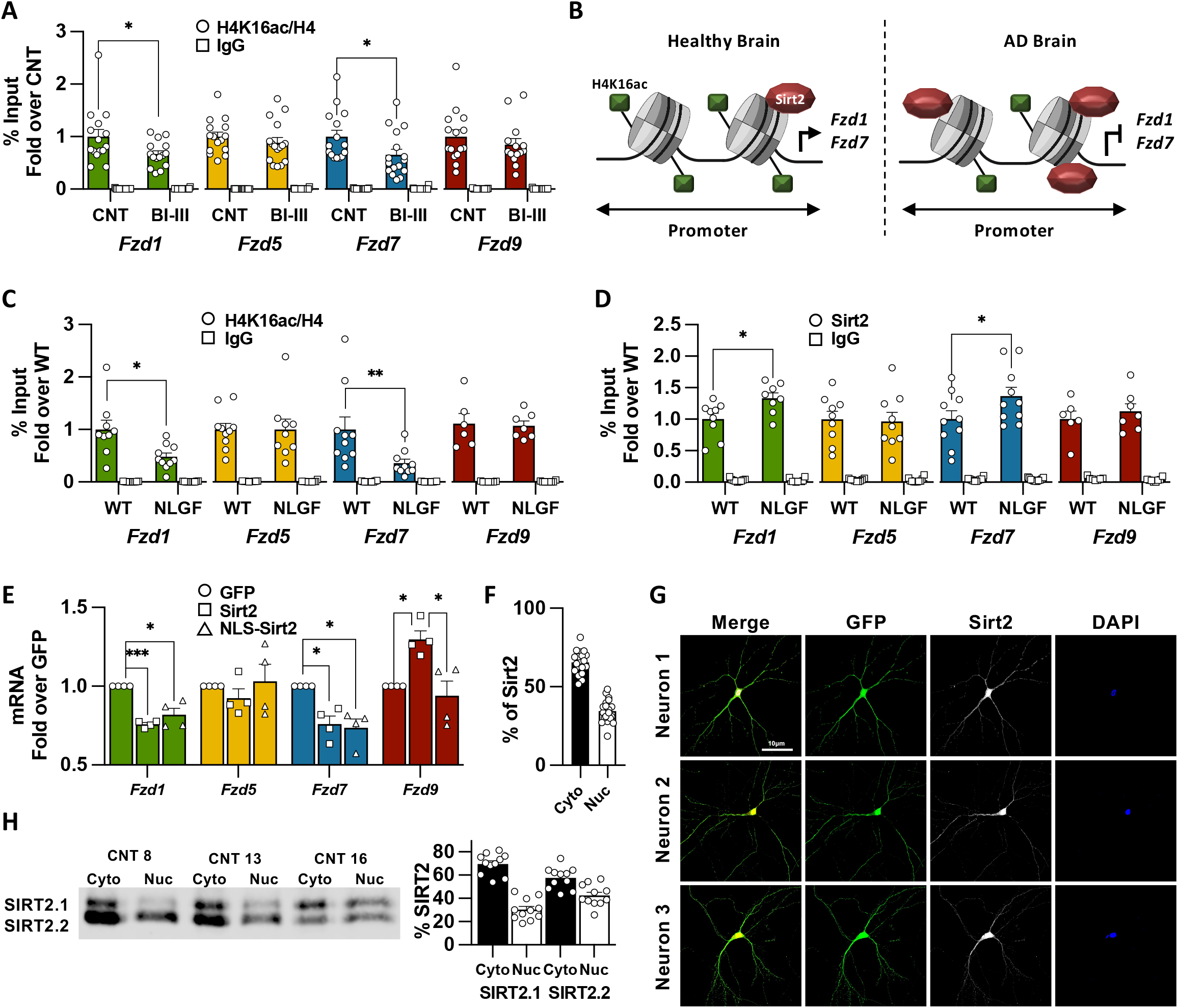
Downregulation of *Fzds* in AD correlates with reduced levels of H4K16ac and the concomitant increase in Sirt2 at their promoters. A) ChIP-qPCR analyses of H4K16ac at the prompters of *FZD1, FZD7, FZD5* and *FZD9* in Human Control and Braak I-III subjects (BI-III) showing reduced acetylation levels at *FZD1* and *FZD7* promoters in AD. H4K16ac levels remain unchanged at *FZD5* and *FZD9* promoters. B) Scheme representing the epigenetic changes observed in AD, where *Fzd1* and *Fzd7* promoters present reduced levels of H4K16ac and increased levels of the histone deacetylase Sirt2. C) ChIP-qPCR experiments showing reduced H4K16ac at *Fzd1* and *Fz7* promoters in NLGF hippocampal samples. No changes are observed for *Fzd5* or *Fzd9*. D) ChIP-qPCR analyses of Sirt2 at the prompters of *Fzd1, Fzd7, Fzd5* and *Fzd9* in WT and NLGF hippocampal samples showing increased Sirt2 levels at *Fzd1* and *Fzd7* promoters in AD. No differences are observed at *Fzd5* or *Fzd9* promoters. E) qPCR analysis showing reduced mRNA levels of *Fzd1* and *Fzd7* in neuronal cultures overexpressing WT Sirt2 or NLS-Sirt2. No changes are observed for *Fzd5*. However, WT Sirt2 induced *Fzd9* transcription. F-G) Quantification (F) and representative images (G) showing Sirt2 is found in the nucleus of postmitotic neurons. In G the first column shows merged images with DAPI (blue), GFP (green) and Sirt2 (White), second column shows GFP, third column shows Sirt2 (white) and last column shows DAPI (blue). H) Quantification of cytosolic and nuclear Sirt2 and representative WB showing that 30-42% of Sirt2 is found in the nucleus in human brain. Data are represented as mean + SEM. Statistical analyses by *t-*Test in A for *FZD1, FZD7* and by Mann-Whitney for *FZD5* and *FZD9*; in C *t-*Test for *Fzd1, Fzd5* and *Fzd9* and by Mann-Whitney for *Fzd7*; in D *t-*Test for all genes; in E one-way ANOVA followed by Games-Howell multiple comparison for all genes. N are indicated in each bar by the number of symbols. Asterisks indicate \**p*<0.05; \*\**p*<0.01, \*\*\**p*<0.005.

Finally, we interrogated which of the three H4K16ac deacetylases (Histone Deacetylases 2 (HDAC), Sirt1 or Sirt2 [27–29]) could be involved in regulating *Fzd1* and *Fzd7* in AD. Interestingly, HDAC2 and Sirt2 play a neurodegenerative role, whereas Sirt1 is neuroprotective [30, 31]. Therefore, we analysed HDAC2 and Sirt2 occupancy at *Fzd* promoters. First, we found that HDAC2 or Sirt2 were not enriched at Fzds promoters in WT (Fig. S1I-J). Interestingly, ChIP-qPCR experiments showed increased Sirt2 occupancy only at *Fzd1* and *Fzd7* promoters in the hippocampus of AD mice (Fig. 2D, S1K). In contrast, no changes were found for HDAC2 levels across all the genes analysed (Fig. S1L). These results show that reduced expression of *Fzd1* and *Fzd7* correlates with reduced levels of H4K16ac and with a concomitant increase of its histone deacetylase Sirt2 at their promoters.

### Nuclear Sirt2 is sufficient to downregulate expression of *Fzds*

To study the possible role of Sirt2 in regulating *Fzds*, we overexpressed Sirt2 in primary neuronal cultures and evaluated *Fzds* mRNA levels. Our results showed that increased Sirt2 expression downregulated *Fzd1* and *Fzd7* expression in neurons, without affecting *Fzd5* and leading to increased *Fzd9* expression (Fig. 2E). Intriguingly, Sirt2 is known to be cytosolic in HEK cells [29](Fig.S2A), whereas we observed a nuclear effect of Sirt2. Interestingly, immunostaining experiments showed that 34.51% of Sirt2 is found in the nucleus in neurons (Fig 2F-G). In addition, 30-42% of Sirt2 is found in nuclear fractions of human hippocampal samples (Fig. 2H, S2D), suggesting that Sirt2 nuclear localisation could be different in postmitotic cells compared to HEK. To drive Sirt2 nuclear translocation, we incorporated a nuclear localisation signal to the Sirt2 N-terminus (NLS-Sirt2; Fig. S2B-C) and studied its impact on *Fzds* expression. We found that NLS-Sirt2 downregulated *Fzd1* and *Fzd7* expression to the same levels of WT Sirt2 in neuronal cultures (Fig. 2E). These results suggest that nuclear Sirt2 is sufficient to downregulate *Fzd1* and *Fzd7* and that Sirt2 nuclear localisation is cell-type dependent.

### Sirt2 inhibition prevents synapse loss and rescues *Fzds* epigenome and transcription in AD

To test if Sirt2 is required for *Fzd1* and *Fzd7* downregulation in AD, we established an AD cellular model: hippocampal primary neurons were cultured for 15DIV and treated overnight with Aβo (Fig. 3A, S2E), leading to *Fzd1* and *Fzd7* reduced expression, reduced H4K16ac and increased Sirt2 levels at their promoters and also synapse loss, without modulating total Sirt2 mRNA or protein levels (Fig. S2F-K). Next, we studied whether Sirt2 inhibition could prevent *Fzd* downregulation and synapse loss. We used a non-toxic concentration of the specific Sirt2 inhibitor AGK2 (Fig. S2L) [32] that leads to increased acetylation of the Sirt2 substrate H3K18ac [33] (Fig S2M). We found that Sirt2 inhibition indeed prevented *Fzd1* and *Fzd7* downregulation in (Fig 3B) and also prevented Aβ-induced synapse loss in our AD cellular model (Fig. 3C). These results suggests that Sirt2 is required for *Fzd1* and *Fzd7* downregulation and synapse loss upon Aβ insult in neurons.

**Figure 3:**
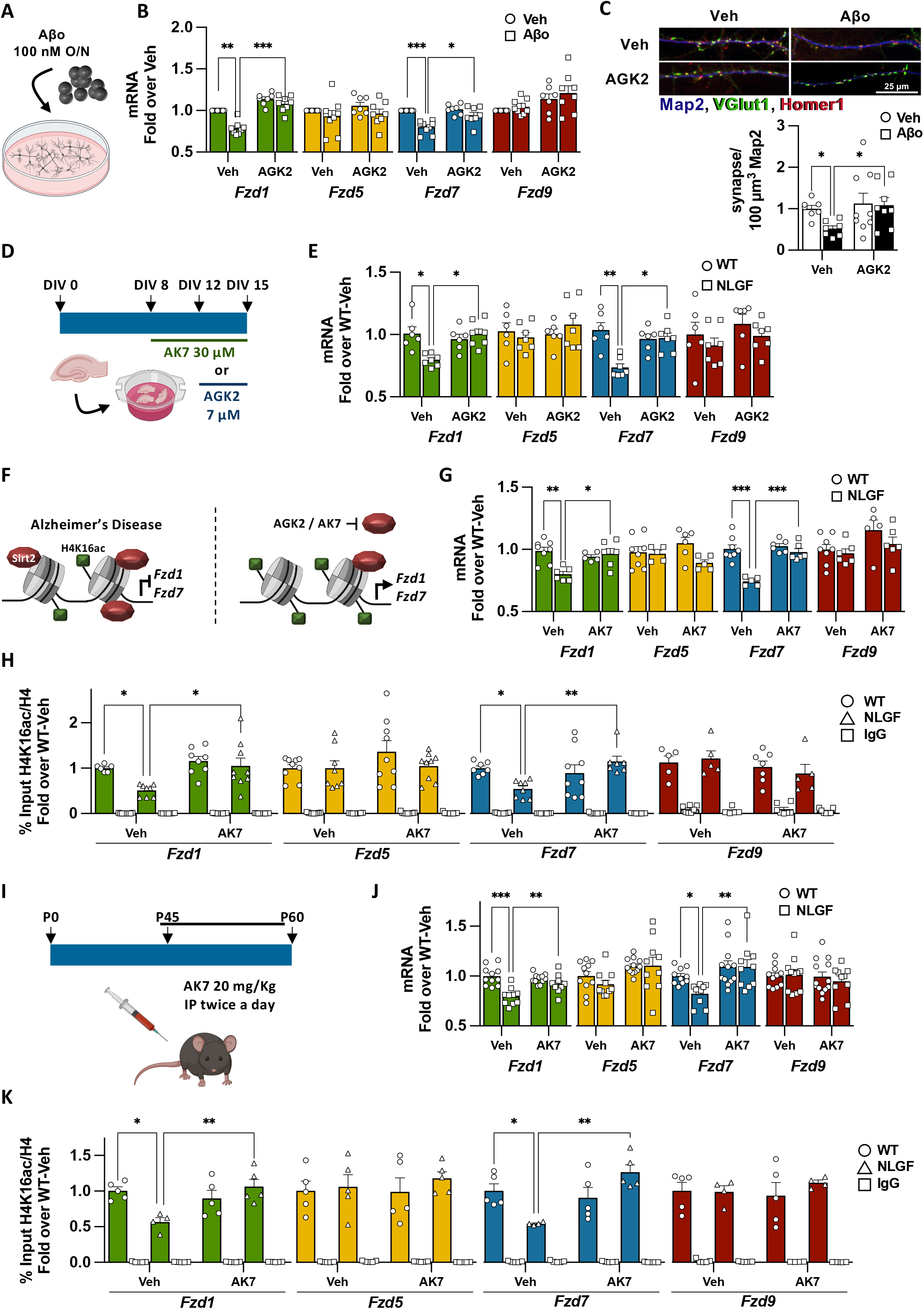
Sirt2 inhibition rescues *Fzd1* and *Fzd7* epigenome and their transcription in AD. A) Scheme representing our cellular AD model, where 15 DIV neuronal cultures were challenged with 100 nM Aβo O/N. B) qPCR analyses of Fzds expression upon Sirt2 inhibition by AGK2 in vehicle (Veh) and Aβo treated neurons, showing that AGK2 prevents *Fzd1* and *Fzd7* downregulation without modulating *Fzd5* or *Fzd9* mRNA levels. C) Representative image and synapse quantification (presynaptic marker vGlut1 (green) on the postsynaptic marker Homer1 (red) and Map2 (blue)) in neuronal cultures treated with the Sirt2 inhibitor AGK2 and challenged with Aβo. Our results show that Sirt2 inhibition prevents Aβo-induced synapse loss. D) Scheme representing AGK2 and AK7 treatments in the *in vitro* AD organotypic model. Hippocampal slices were cultured for 15 DIV and treated with AGK2 for 72 hours or with AK7 for 7 days. E) qPCR analyses of total mRNA levels from WT and NLGF hippocampal cultures treated with vehicle or AGK2. Our results show that AGK2 treatment rescues *Fzd1* and *Fzd7* mRNA levels without modulating *Fzd5* or *Fzd9* mRNA levels. F) Scheme representing the epigenetic state of *Fzd1* and *Fzd7* promoters in AD and how Sirt2 inhibition rescues H4K16ac levels and their mRNA expression. G) qPCR results show that AK7 treatment rescues *Fzd1* and *Fzd7* mRNA levels in AD treated cultures, without modulating *Fzd5* or *Fzd9* mRNA levels. H) ChIP-qPCR showing that AK7 treatment rescues the levels of H4K16ac at *Fzd1* and *Fzd7* promoters in hippocampal organotypic cultures of NLGF while not changing the levels of this pro-transcriptional histone mark in WT or at *Fzd5* and *Fzd9* promoter. I) Scheme showing the dosage regime for *in vivo* inhibition of Sirt2 by intraperitoneal injections of 20 mg/Kg of AK7 twice a day for 15 days, from 1.5 to two months old animals. J) qPCR analyses of total mRNA levels from WT and NLGF hippocampal samples treated with vehicle or AK7. Our results show that AK7 treatment rescues *Fzd1* and *Fzd7* mRNA levels back to WT in NLGF treated animals and does not show any effect on *Fzd5* or *Fzd9* mRNA levels. K) ChIP-qPCR showing that AK7 treatment rescues the levels of H4K16ac at *Fzd1* and *Fzd7* promoters in AD while not changing are observed in WT or at *Fzd5* and *Fzd9* promoters. Data are represented as mean + SEM. Statistical analyses by Two-way ANOVA followed by Games-Howell post hoc in B for all genes analysed; in C by Kruskal-Wallis followed by Dunn’s multiple comparison; in E by Two-way ANOVA followed by Tukey’s post hoc for *Fzd1, Fzd5* and *Fzd7* and by Kruskal-Wallis followed by Dunn’s multiple comparison for *Fzd9*; in G Two-way ANOVA followed by Tukey’s post hoc for *Fzd1, Fzd7* and *Fzd9* and by Kruskal-Wallis followed by Dunn’s multiple comparison for *Fzd5*; in H Kruskal-Wallis followed by Dunn’s multiple comparison for all *Fzd1, Fzd5* and *Fzd7* and by Two-way ANOVA followed by Tukey’s post hoc for *Fzd9*; in G Two-way ANOVA followed by Tukey’s post hoc for *Fzd1* and *Fzd9* and Kruskal-Wallis followed by Dunn’s multiple comparison for *Fzd5* and *Fzd7*; in K Two-way ANOVA followed by Tukey’s post hoc for *Fzd1* and *Fzd5* and Two-way ANOVA followed by Games-Howell post hoc for *Fzd7* and *Fzd9*. N are indicated in each bar by the number of symbols. Asterisks indicate \**p*<0.05; \*\**p*<0.01; \*\*\**p*<0.005.

To further study the role of Sirt2 in regulating Fzds expression in the context of AD, we prepared hippocampal organotypic cultures (HOC) from WT and NLGF animals. Consistent with our results in BI-III and NLGF mice, we found reduced H4K16ac and increased Sirt2 levels at *Fzd1* and *Fzd7* promoters and a concomitant reduction in their transcription in NLGF-HOC (Fig. S3A-D). Next, we studied whether Sirt2 inhibition could rescue *Fzds* expression in our HOC AD model. First, the Sirt2 specific inhibitor AGK2 showed no toxicity (Fig. 3D, S3E, Table S2) and effectively supressed Sirt2 activity as shown by increased acetylation of the Sirt2 substrate H3K56ac [33] (Fig. S3F). Indeed, Sirt2 inhibition by AGK2 rescued *Fzd1* and *Fzd7* expression in the NLGF-HOC, without affecting control genes (Fig. 3E-F). Second, we treated our HOC model with a second specific and structurally distinct Sirt2 inhibitor; AK7 (Table S2) [32]. AK7 treatment was not toxic and effectively supressed Sirt2 activity (Fig. 3D, S3E, S3G). Importantly, Sirt2 inhibition by AK7 rescued *Fzd1* and *Fzd7* mRNA levels in the NLGF-HOC model, without modulating control genes (Fig. 3G). Thus, Sirt2 inhibition by two distinct small molecules suggest that Sirt2 represses *Fzd1* and *Fzd7* in the context of AD. Finally, we analysed whether AK7 treatment also rescued H4K16ac in NLGF-HOC. Indeed, we found that the levels of this pro-transcriptional histone mark were restored at *Fzd1* and *Fzd7* promoters in the NLGF-HOC treated cultures (Fig. 3H, S3H). Interestingly, Sirt2 has other histone substrates, including H3K18ac and H3K56ac [33]. However, we found that these two marks were not enriched at *Fzd1* or *Fzd7* promoters in WT (Fig. S3I-J), and no differences were observed in hippocampus of the NLGF model compared to control (Fig. S3K-L). These results suggest that Sirt2 impairs *Fzd1* and *Fzd7* transcription by specifically reducing H4K16ac levels in their promoters in the AD context.

Finally, we tested the role of Sirt2 in regulating *Fzd1* and *Fzd7* transcription *in vivo* by using the Sirt2 inhibitor AK7, which crosses the blood brain barrier [34]. Mice were injected intraperitoneally with 20 mg/Kg twice a day for 15 days (Fig. 3I), as previously reported [35]. AK7 administration effectively inhibited Sirt2 in the brain (Fig. S3M), rescuing *Fzd1* and *Fzd7* expression (Fig. 3J) and H4K16ac levels at their promoters (Fig. 3H) in NLGF animals. Similar to our *in vitro* studies, AK7 did not modulate the mRNA levels of control genes or the levels of H4K16ac at their promoters (Fig. 3K, S3N). Interestingly, we found no changes in Aβ_42_ levels (Fig. S3O), as previously reported with the same AK7 dosage in two different AD models [35]. Collectively, these results show that *Fzd1* and *Fzd7* genes are repressed through Sirt2 deacetylation of H4K16ac in the context of AD (Fig. 3F).

### FoxO1 recruits Sirt2 to *Fzd1* and *Fzd7* promoters in AD

Increased Sirt2 occupancy at *Fzd1* and *Fzd7* promoters suggests that Sirt2 levels might be upregulated in AD. To test this hypothesis, we analysed SIRT2 mRNA and protein levels in BI-III human hippocampal samples and found no changes (Fig. S4A-B). Similarly, no changes in Sirt2 protein levels were observed in NLGF-HOC model (Fig. S4C), but we observed reduced *Sirt2* mRNA levels in NLGF-HOC (Fig. S4D). In addition, no differences in SIRT2 nuclear levels were observed in BI-III, but we found increased nuclear Sirt2 in NLGF-HOC (Fig. 4A, S4E-F). These results indicate that increased Sirt2 occupancy at *Fzd1* and *Fzd7* promoters does not correlate with increased total/nuclear levels of Sirt2 in the human BI-III, suggesting that Sirt2 might be recruited to Fzds promoters by co-factors.

**Figure 4:**
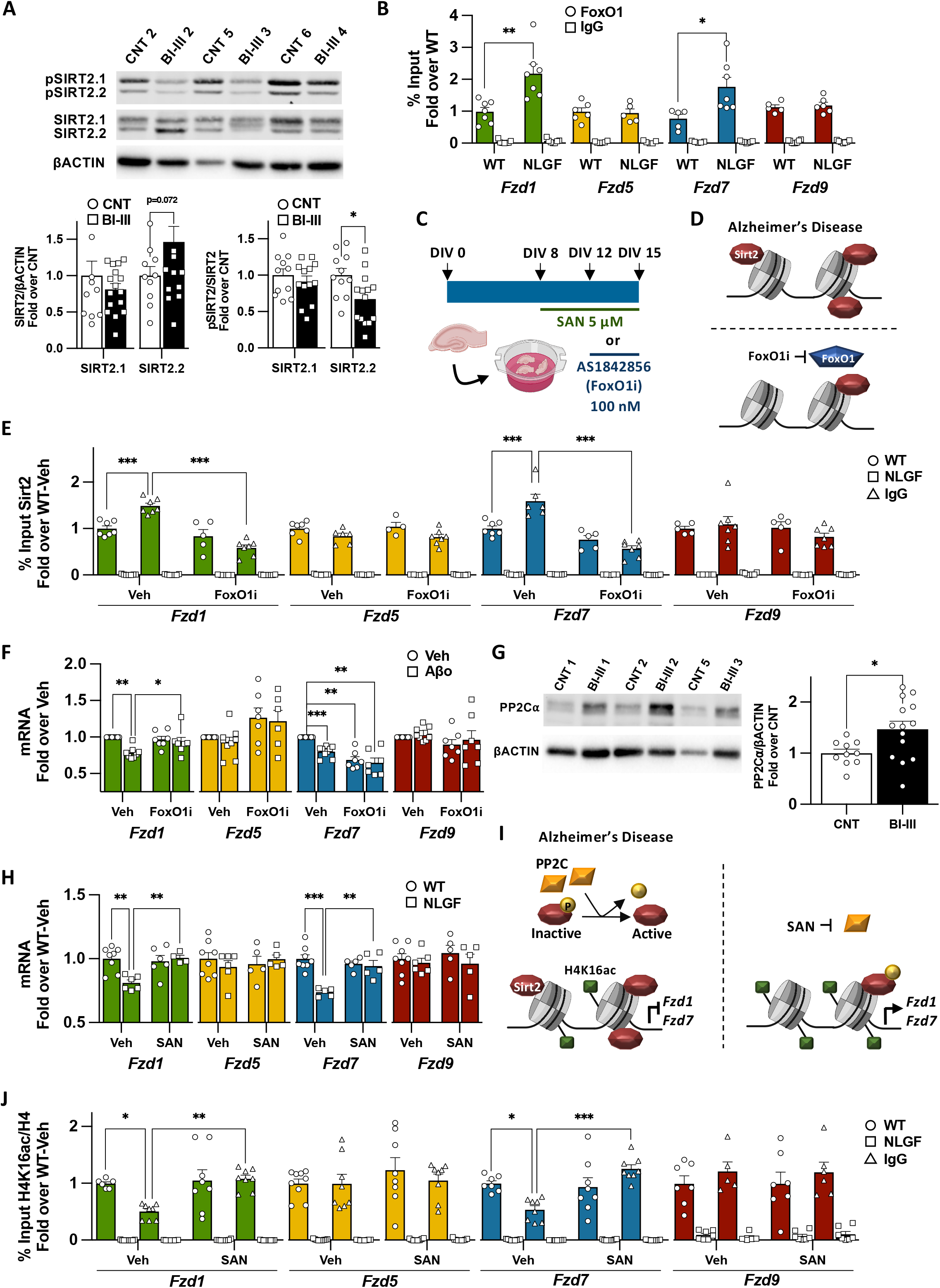
Increased Sirt2 activity in AD impairs Fzds transcription. A) WB analyses of total and pSirt2 levels in hippocampal nuclear extracts of human control/BI-III subjects showing decreased levels of pSirt2.2 at early AD. B) ChIP-qPCR analyses of FoxO1 in WT and NLGF hippocampal samples showing increased FoxO1 levels at *Fzd1* and *Fzd7* promoters in AD. No differences are observed at *Fzd5* or *Fzd9* promoters. C) Scheme representing Sanguinarine (SAN) and AS1842856 FoxO1 inhibitor (Fox1Oi) treatment in the *in vitro* AD organotypic model for and 7 days and 72h respectively. D) Scheme representing the levels of Sirt2 at *Fzd1* and *Fzd7* and upon FoxO1 inhibition in AD. E) ChIP-qPCR showing that FoxO1i treatment reduces Sirt2 levels at *Fzd1* and *Fzd7* promoters in hippocampal organotypic cultures of NLGF while not changing the levels of Sirt2 in WT or at *Fzd5* or *Fzd9* promoter, suggesting FoxO1 recruits Sirt2 to *Fzd1* and *Fzd7* promoters in AD. F) qPCR analyses of Fzds expression upon FoxO1 inhibition in vehicle (Veh) and Aβo treated neurons, showing that FoxO1i prevents *Fzd1* downregulation without modulating *Fzd5* or *Fzd9* mRNA levels. FoxO1 inhibition downregulates *Fzd7* expression *per se* and fails to prevent its downregulation in Aβo treated neurons. G) WB analyses of PP2C⍰in hippocampal nuclear extracts of human control/BI-III subjects, sowing increased levels of PP2C⍰in human BI-III group. H) qPCR analyses of total mRNA levels from WT and NLGF hippocampal organotypic cultures treated with vehicle or SAN. Our results show that SAN treatment rescues *Fzd1* and *Fzd7* mRNA levels and does not show any effect on *Fzd5* or *Fzd9* mRNA levels. I) Scheme representing increased nuclear levels of the phosphatase PP2C in AD and how PP2C inhibition in AD rescues H4K16ac levels at Fzd1 and Fzd7 promoters and their transcription. J) ChIP-qPCR showing that SAN treatment rescues the levels of H4K16ac at *Fzd1* and *Fzd7* promoters in AD while not changes are observed in WT or at *Fzd5* and *Fzd9* promoter. Data are represented as mean + SEM. Statistical analyses by *t-*Test in A for total Sirt2.2 and nuclear Sirt2.1 and Srit2.2, and by Mann-Whitney for total Sirt2.1; in B in by *t-*Test for all genes analysed; E Two-way ANOVA followed by Tukey’s post hoc for all genes analysed; in F by Two-way ANOVA followed by Games-Howell post hoc for all genes analysed; in G by *t-*Test; in H Two-way ANOVA followed by Tukey’s post hoc for *Fzd1, Fzd7* and *Fzd9* and Kruskal-Wallis followed by Dunn’s multiple comparison for *Fzd5*; in J Two-way ANOVA followed by Tukey’s post hoc for *Fzd7* and *Fzd9* and Kruskal-Wallis followed by Dunn’s multiple comparison for *Fzd1* and *Fzd5*. N are indicated in each bar by the number of symbols. Asterisks indicate \**p*<0.05; \*\**p*<0.01; \*\*\**p*<0.005.

Sirt2 interacts with FoxO1 and FoxO3a transcription factors [36, 37], which could recruit Sirt2 to specific loci. Using CiiiDER [38], we found putative FoxO1, but not FoxO3a, binding sites at *Fzd1* and *Fzd7* promoters (Fig. S4G). Next, we analysed FoxO1 occupancy at *Fzd1* and *Fzd7* promoters in AD. We found increased FoxO1 levels at *Fzd1* and *Fzd7* promoters in NLGF hippocampal samples (Fig. 4B, S4H), suggesting that FoxO1 could contribute to the recruitment of Sirt2 to *Fzd1* and *Fzd7* promoters in the context of AD. To test this hypothesis, we treated HOC with the specific FoxO1 activity inhibitor AS1842856 (FoxO1i, Fig. 4C, Table S2) [39], and found no cytotoxicity (Fig. S4I). We next analysed Sirt2 occupancy upon FoxO1 inhibition and found reduced Sirt2 levels at *Fzd1* and *Fzd7* promoters in the NLGF-HOC model (Fig. 4D-E). No changes were observed at control gene promoters (Fig. 4E, S4J). However, we found reduced Sirt2 levels at *Hoxa1* promoter, which has three putative FoxO1 binding sites (Fig. S4G, S4J).

To further test the role of FoxO1 in repressing *Fzd1* and *Fzd7* in the context AD, we treated our cellular AD model with a non-toxic concentration of FoxO1 inhibitor (Fig. S4K), which indeed prevented Aβ-induced *Fzd1* downregulation (Fig. 4F). No changes were observed for control genes (Fig. 4F). However, FoxO1 inhibition downregulated *Fzd7* expression in WT samples and consequently failed to prevent *Fzd7* downregulation in the context of AD (Fig. 4F). Interestingly, *Fzd7* was the only Fzd receptor that displayed high FoxO1 occupancy in WT (Fig. S4L). Furthermore, FoxO1 inhibition in WT-HOC led to reduced H4K16ac levels at *Fzd7* promoter (Fig. S4M). Together these results suggest that FoxO1 activity is required for *Fzd7* basal expression.

Next, we treated neurons with the Sirt2 inhibitor AGK2 together with the FoxO1 inhibitor. Our results showed that inhibition of these two proteins prevented *Fzd1* downregulation in the context of AD (Fig. S4N) with no changes in control genes (Fig. S4N). In contrast, this double inhibition led to *Fzd7* downregulation as we observed with inhibition of FoxO1 alone (Fig. S4N). Altogether, these results suggest that FoxO1 recruits Sirt2 to *Fzd1* and *Fzd7* promoters leading to their downregulation in the context of AD. The difference in the response to the FoxO1 inhibition between *Fzd1* and *Fzd7* might reflect difference in the regulation under basal conditions.

### Increased nuclear Sirt2 activity represses *Fzd1* and *Fzd7* in AD

Sirt2 activity can be modulated by phosphorylation. We therefore analysed the phosphorylation of Sirt2 at its Serine 331 (pSirt2) which inhibits its activity [40, 41]. We found reduced pSIRT2 levels in nuclear fractions of BI-III hippocampal samples (Fig. 4A), without changes in total pSIRT2 levels (Fig. S4A). Similarly, lower levels of nuclear and total pSirt2 were observed in NLGF-HOC (Fig. S4C, S4F). Decreased levels of Sirt2 inhibitory phosphorylation could be regulated by specific phosphatases, such as the Sirt2 phosphatase *PP2C*⍰[41], which is upregulated at the RNA level in an AD model [42]. We therefore analysed the expression of the Sirt2 phosphatases PP2C⍰/β [41] and found no changes in the mRNA or total protein levels in hippocampal samples from human BI-III or NLGF-HOC (Fig. S5A-F). Next, we analysed nuclear localisation and found increased levels of PP2C⍰, but not PP2Cβ, in BI-III subjects (Fig. 4G, S5G). Importantly, Pp2c⍰was also upregulated in nuclear samples of NLGF-HOC (Fig S5H). These results suggest that increased nuclear levels of PP2C⍰could lead to Sirt2 nuclear hyperactivity, favouring the repression of *Fzd1* and *Fzd7* in AD.

To establish the role of PP2C in *Fzd1* and *Fzd7* expression in AD, HOC were treated with a non-toxic concentration of sanguinarine (SAN), a PP2C specific inhibitor [43] (Fig. 4D, S4I, Table. S2) and we analysed the impact of SAN on the expression of *Fzd1* and *Fzd7*. Our results showed that Pp2c inhibition rescued *Fzd1* and *Fzd7* expression and H4K16ac levels in the NLGF-HOCs, without affecting the expression of control genes (Fig. 4H-J, S5I). Consistently, increased pSirt2 levels were only found in NLGF-HOC SAN-treated samples (Fig. S5J), supporting the idea that increased nuclear Pp2c levels are responsible for hyperactive Sirt2. Finally, we inhibited Pp2c and Sirt2 in HOC and found that co-inhibiting both enzymes also rescued *Fzd1* and *Fzd7* expressing in NLGF-HOC (Fig. S5K). These results suggest that Pp2c is upstream of Sirt2 in repressing *Fzd1* and *Fzd7* in the context of AD.

## DISCUSSION

In this study, we present novel findings demonstrating that *Fzd1* and *Fzd7* genes are epigenetically repressed by hyperactive nuclear Sirt2 in AD. In addition, Sirt2 recruitment to *Fzd1* and *Fzd7* promoters depends on FoxO1 activity, leading to increased Sirt2 levels at these promoters and a concomitant H4K16ac deacetylation resulting in the repression of *Fzd1* and *Fzd7* genes in AD (Fig. 5).

**Figure 5:**
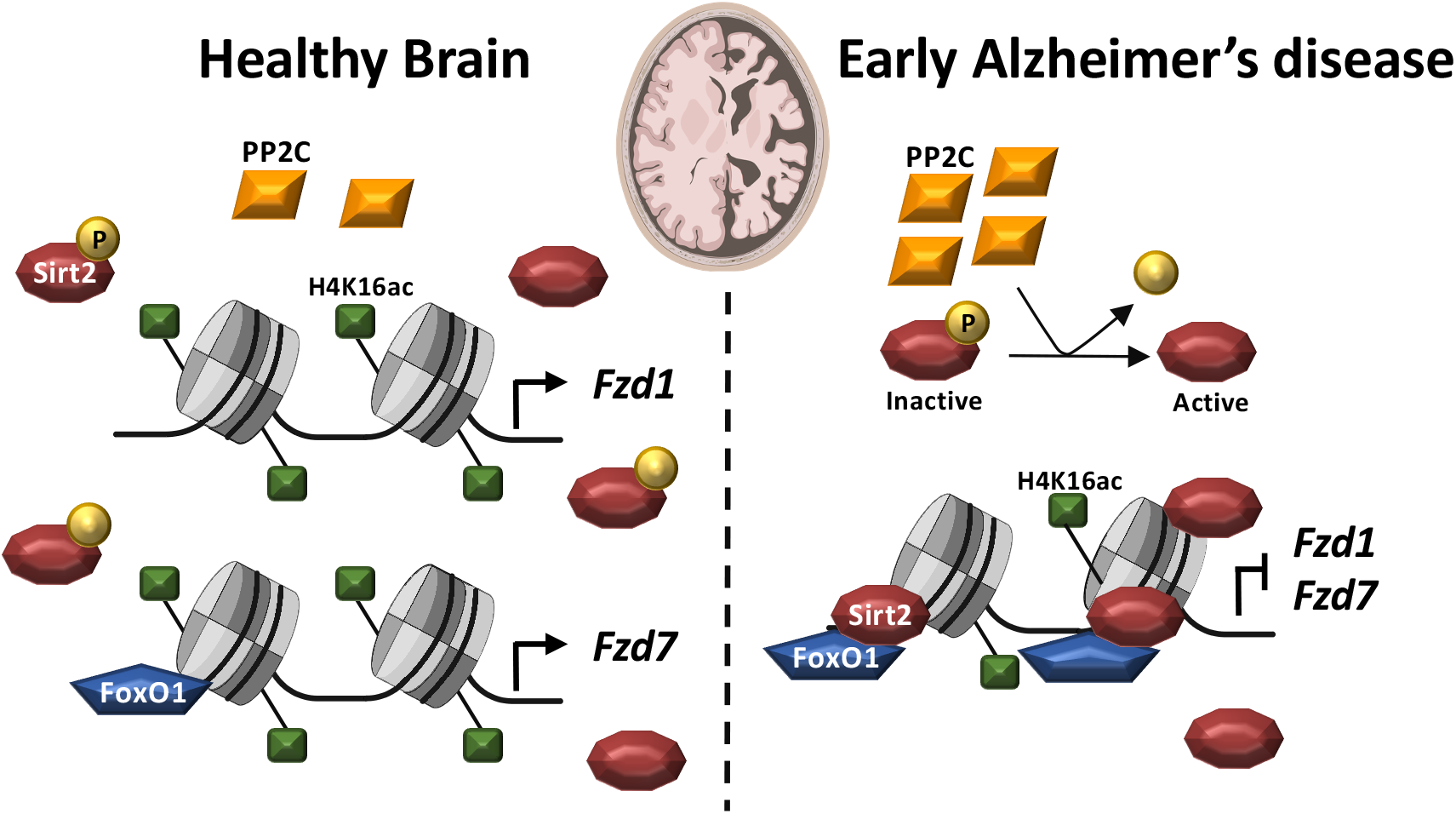
Schematic model of *Fzd1* and *Fzd7* regulation by Sirt2 in AD. Scheme representing the nuclear localization of the histone deacetylase Sirt2, its phosphatase PP2C and the epigenetic regulation of *Fzd1* and *Fzd7* in the healthy brain and in AD. In the healthy brain, high levels of H4K16ac and low levels of Sirt2 coexist at *Fzd1* and *Fzd7* prompters. In addition, high levels of FoxO1 are present at *Fzd7* promoter, altogether leading to *Fzd1* and *Fzd7* transcription. In AD, increased nuclear levels of the phosphatase PP2C activates Sirt2 by removing its inhibitory phosphorylation. In turn, FoxO1 recruits Sirt2 to *Fzd1* and *Fzd7* promoters leading to reduced levels of H4K16ac and impairing *Fzd1* and *Fzd7* transcription.

Wnt signalling has been linked to AD by the identification of three LRP6 genetic variants and by the synaptotoxic role of the Wnt antagonist Dkk1, which is required for Aβ-mediated synapse loss [8–11]. Here, we report for the first time that two Fzds with synaptic function display reduced expression in the hippocampus of BI-III patients and the NLGF model. Our results showed a reduction in neuronal *Fzd1* and *Fzd7* expression in AD, which seems to be hippocampus specific as this was not observed in other brain regions in the human condition at early disease stages [44, 45] (Fig S5L). Consistently, we observed *Fzd1* and Fzd7 downregulation in the hippocampus of the overexpressing AD model APP/PS1 (Fig. S5M), but not in J20 (Fig. S5N) or other models [46], at a similar disease stage [47, 48]. Further studies are required to reconcile these results. Nonetheless, we observed *Fzd1* and *Fzd7* downregulation by Aβo in hippocampal neuronal cultures, in NLGF-HOC, *in vivo* in the NLGF and APP/PS1 hippocampus and more importantly in the human hippocampus of BI-III subjects.

Reduced Fzd1 and Fzd7 expression in AD could impact both sides of the synapse as these proteins are localised at the pre- and post-synaptic side respectively [17, 18]. Postsynaptically, Fzd7 is required for dendritic arborization during postnatal development [49], spine formation and growth and also for LTP [17]. At the presynaptic site, Fzd1 is sufficient and required for presynaptic assembly[18]. Interestingly, Wnt3a prevents Aβ-induced synapse loss in a Fzd1-dependent manner [50]. We observe reduced *Fzd1* expression in AD, which could contribute to synapse vulnerability in this condition. Together, these results suggest that reduced levels of Fzd1 and Fzd7 could lead to impaired synaptic plasticity and synapse loss at early stages of AD.

*Fzd1* and *Fzd7* expression are regulated by epigenetic mechanisms such as noncoding RNAs and DNA methylation in different biological processes and diseases [51–54]. In this study, we showed a novel role for Sirt2 in repressing *Fzd1* and *Fzd7* by specifically deacetylating the histone H4K16ac at their promoters in the context of AD (Fig. 5), whereas other Sirt2 histone substrates remain unchanged. Interestingly, H4K16ac has been linked to Fzds regulation in brain development: H4K16ac deacetylation by Sirt1 regulates *Fzd5* and *Fzd7* transcription during cortical neurogenesis [55]. Together these results strongly suggest that H4K16ac deacetylation regulates the transcription of different *Fzds* in development and in AD. These studies also show different roles for Sirt1 and Sirt2, consistent with the idea that Sirt1 is neuroprotective whereas Sirt2 plays a neurodegenerative role [30, 31].

Increasing evidence suggests a neurodegenerative role for Sirt2. First, a genetic variant of *SIRT2* is linked to LOAD in *APOE*_ε_*4*-negative population [56]. Second, *in vivo* Sirt2 inhibition improves cognition in three AD models [35, 57]. Third, increased SIRT2 levels are observed in a cellular AD model [58, 59]. However, our results show no changes in SIRT2 levels in human hippocampal BI-III samples. Fourth, Sirt2 inhibition leads to reduced Aβ levels in the APP/PS1 AD model when dosed with 100 mg/Kg of AK7 for three weeks [57]. However, we found no changes in Aβ levels upon Sirt2 inhibition when dosing animals with 20 mg/Kg of AK7 for two weeks. Our results are in line with a previous report showing no changes in Aβ levels in two AD models treated with the same AK7 regime [35]. These apparently contradictory results suggest that shorter Sirt2 inhibition using a low AK7 dosage is sufficient to rescue molecular and memory deficits in AD models independently from Aβ levels. In contrast, longer Sirt2 inhibition with higher AK7 dosage also reduces Aβ levels. Interestingly, we observed a specific reduction of Sirt2 phosphorylation in nuclear fractions of BI-III patients, a post-translational modification known to inhibit Sirt2 deacetylase activity [40, 41]. These results suggest the presence of hyperactive nuclear Sirt2 in AD. We also showed increased nuclear levels of PP2C⍰, a Sirt2 phosphatase, in BI-III subjects, in line with previous results showing increased mRNA levels of PP2C⍰in an AD mouse model [42]. Consistently, we found that PP2C-induced nuclear Sirt2 hyperactivity is upstream of H4K16 deacetylation by Sirt2 at *Fzd1* and *Fzd7* promoter in the context of AD (Fig. 5).

Increased Sirt2 levels at *Fzd1* and *Fzd7* promoters suggest that Sirt2 is specifically recruited by a co-factor with DNA binding capacity. Interestingly, Sirt2 interacts with the transcription factor FoxO1 [36], which has predicted binding sites at *Fzd1* and *Fzd7* promoter region. FoxO1 can positively or negatively regulate gene transcription in different biological conditions [60]. We found that FoxO1 is enriched at *Fzd7* and is required for its basal transcription in neurons as FoxO1 inhibition downregulated *Fzd7* expression and reduced the levels of H4K16ac at its promoter under basal conditions. But, FoxO1 did not modulate *Fzd1* expression under basal conditions. In contrast, our results showed that FoxO1 inhibition prevents Sirt2 recruitment to *Fzd1* and *Fzd7* promoters and prevents *Fzd1* downregulation in the context of AD, suggesting that FoxO1 acts as a co-repressor. Together, these results suggest that FoxO1 acts as a repressor for *Fzd1* in AD and that this transcription factor has a dual role for *Fzd7*: from positive regulation of *Fzd7* transcription in basal conditions to negative regulation of *Fzd7* transcription in AD context. Interestingly, this transcriptional repression could also regulate other genes with synaptogenic or neuroprotective attributes such as the Wnt ligands *Wnt3a, Wnt5a/b* or the neurotropic factors *Ngf* or *Ntf3* [13, 50, 61, 62], as all of them present putative FoxO1 binding sites in their promoters when analysed by CiiiDER (Fig. S5O). This mechanism could also regulate other genes implicated in AD. This epigenetic regulation of Wnt receptors by the Sirt2-H4K16ac could also modulate the expression of these genes in other cellular contexts and in diseases.

In summary, we report a novel role for nuclear Sirt2 in regulating *Fzd* receptors in AD. We propose that nuclear Sirt2 is hyperactivated in AD, and that FoxO1 recruits Sirt2 to *Fzd1* and *Fzd7* promoters leading to reduced H4K16ac, which in turn impairs their transcription. Thus, Sirt2 is a promising target for developing new AD therapies to restore the expression of key Wnt receptors.

## SUPPLEMENTARY INFORMATION

For further information see Supplementary Figures, Supplementary Methods and Supplementary Tables.

## AUTHOR’S CONTRIBUTION

E.P. and P.C.S. conceived this study. E.P., N.M.F., M.P. and K.V. performed *in vivo* experiments. E.P., S.J. and P.W. performed smFISH and experiments in human samples. E.P., P.P-V. and S.B. performed *in vitro* experiments. E.P. and S.T. performed cell biology experiments. T.S. and T.C.S. provided the NLGF line. E.P. and P.C.S. wrote the manuscript with input from all authors.

## ACKNOWLEDGMENTS/FUNDINGS

We thank Prof Jose A. Esteban for kindly providing us with APP/PS1 hippocampal samples, Dr William Andrews for his help in cloning and the members of the Salinas Lab for their input on the project and manuscript. This work was funded by the MRC (P.C.S.: MR/M024083/1), Alzheimer’s Society (P.C.S.: AS-PG-18-008), Alzheimer’s Research UK (P.C.S. and E.P.: ARUK-PG2018A-002; P.W.: ARUK-2018DDI-UCL), European Commission Horizon 2020 (E.P.: H2020 MSCA-IF 749209).

## REFERENCES

1. Terry RD, Masliah E, Salmon DP, Butters N, DeTeresa R, Hill R, et al. Physical basis of cognitive alterations in alzheimer’s disease: Synapse loss is the major correlate of cognitive impairment. Ann Neurol. 1991;30:572–580.

2. Colom-Cadena M, Spires-Jones T, Zetterberg H, Blennow K, Caggiano A, DeKosky ST, et al. The clinical promise of biomarkers of synapse damage or loss in Alzheimer’s disease. Alzheimers Res Ther. 2020;12:21.

3. Forner S, Baglietto-Vargas D, Martini AC, Trujillo-Estrada L, LaFerla FM. Synaptic Impairment in Alzheimer’s Disease: A Dysregulated Symphony. Trends Neurosci. 2017;40:347–357.

4. Mucke L, Selkoe DJ. Neurotoxicity of amyloid β-protein: synaptic and network dysfunction. Cold Spring Harb Perspect Med. 2012;2:a006338.

5. Caricasole A, Copani A, Caraci F, Aronica E, Rozemuller AJ, Caruso A, et al. Induction of Dickkopf-1, a Negative Modulator of the Wnt Pathway, Is Associated with Neuronal Degeneration in Alzheimer’s Brain. J Neurosci. 2004;24:6021–6027.

6. Jolly S, Lang V, Koelzer VH, Sala Frigerio C, Magno L, Salinas PC, et al. Single-Cell Quantification of mRNA Expression in The Human Brain. Sci RepoRtS |. 2019;9:12353.

7. Rosi MC, Luccarini I, Grossi C, Fiorentini A, Spillantini MG, Prisco A, et al. Increased Dickkopf-1 expression in transgenic mouse models of neurodegenerative disease. 2010;112:1539–1551.

8. Purro SA, Dickins EM, Salinas PC. The Secreted Wnt Antagonist Dickkopf-1 Is Required for Amyloid -Mediated Synaptic Loss. J Neurosci. 2012;32:3492–3498.

9. Sellers KJ, Elliott C, Jackson J, Ghosh A, Ribe E, Rojo AI, et al. Amyloid β synaptotoxicity is Wnt-PCP dependent and blocked by fasudil. Alzheimers Dement. 2018;14:306–317.

10. De Ferrari G V, Papassotiropoulos A, Biechele T, Wavrant De-Vrieze F, Avila ME, Major MB, et al. Common genetic variation within the low-density lipoprotein receptor-related protein 6 and late-onset Alzheimer’s disease. Proc Natl Acad Sci U S A. 2007;104:9434–9439.

11. Alarcón MA, Medina MA, Hu Q, Avila ME, Bustos BI, Pérez-Palma E, et al. A novel functional low-density lipoprotein receptor-related protein 6 gene alternative splice variant is associated with Alzheimer’s disease. Neurobiol Aging. 2013;34:1709.e9-18.

12. Alvarez AR, Godoy JA, Mullendorff K, Olivares GH, Bronfman M, Inestrosa NC. Wnt-3a overcomes beta-amyloid toxicity in rat hippocampal neurons. Exp Cell Res. 2004;297:186–196.

13. Cerpa W, Farías GG, Godoy JA, Fuenzalida M, Bonansco C, Inestrosa NC. Wnt-5a occludes Abeta oligomer-induced depression of glutamatergic transmission in hippocampal neurons. Mol Neurodegener. 2010;5:3.

14. Marzo A, Galli S, Lopes D, McLeod F, Podpolny M, Segovia-Roldan M, et al. Reversal of synapse degeneration by restoring Wnt signalling in the adult hippocampus. Curr Biol. 2016;26:2551–2561.

15. Killick R, Ribe EM, Al-Shawi R, Malik B, Hooper C, Fernandes C, et al. Clusterin regulates β-amyloid toxicity via Dickkopf-1-driven induction of the wnt-PCP-JNK pathway. Mol Psychiatry. 2014;19:88–98.

16. Nusse R, Clevers H. Wnt/β-Catenin Signaling, Disease, and Emerging Therapeutic Modalities. Cell. 2017;169:985–999.

17. McLeod F, Bossio A, Marzo A, Ciani L, Sibilla S, Hannan S, et al. Wnt Signaling Mediates LTP-Dependent Spine Plasticity and AMPAR Localization through Frizzled-7 Receptors. Cell Rep. 2018;23:1060–1071.

18. Varela-Nallar L, Grabowski CP, Alfaro IE, Alvarez AR, Inestrosa NC. Role of the Wnt receptor Frizzled-1 in presynaptic differentiation and function. Neural Dev. 2009;4:41.

19. Ramírez VT, Ramos-Fernández E, Henríquez JP, Lorenzo A, Inestrosa NC. Wnt-5a/frizzled9 receptor signaling through the Gαo-Gβγ complex regulates dendritic spineformation. J Biol Chem. 2016;291:19092–19107.

20. Sahores M, Gibb A, Salinas PC. Frizzled-5, a receptor for the synaptic organizer Wnt7a, regulates activity-mediated synaptogenesis. Development. 2010;137:2215–2225.

21. Palomer E, Buechler J, Salinas PC. Wnt signaling deregulation in the aging and Alzheimer’s brain. Front Cell Neurosci. 2019;13.

22. Saito T, Matsuba Y, Mihira N, Takano J, Nilsson P, Itohara S, et al. Single App knock-in mouse models of Alzheimer’s disease. Nat Neurosci. 2014;17:661–663.

23. Mathys H, Davila-Velderrain J, Peng Z, Gao F, Mohammadi S, Young JZ, et al. Single-cell transcriptomic analysis of Alzheimer’s disease. Nature. 2019;570:332–337.

24. Zhang Y, Chen K, Sloan SA, Bennett ML, Scholze AR, O’Keeffe S, et al. An RNA-Sequencing Transcriptome and Splicing Database of Glia, Neurons, and Vascular Cells of the Cerebral Cortex. J Neurosci. 2014;34.

25. Horikoshi N, Kumar P, Sharma GG, Chen M, Hunt CR, Westover K, et al. Genome-wide distribution of histone H4 Lysine 16 acetylation sites and their relationship to gene expression. Genome Integr. 2013;4:3.

26. Nativio R, Donahue G, Berson A, Lan Y, Amlie-Wolf A, Tuzer F, et al. Dysregulation of the epigenetic landscape of normal aging in Alzheimer’s disease. Nat Neurosci. 2018;21:497–505.

27. Ma P, Schultz RM. Histone Deacetylase 2 (HDAC2) Regulates Chromosome Segregation and Kinetochore Function via H4K16 Deacetylation during Oocyte Maturation in Mouse. PLoS Genet. 2013;9.

28. Vaquero A, Scher M, Lee D, Erdjument-Bromage H, Tempst P, Reinberg D. Human SirT1 interacts with histone H1 and promotes formation of facultative heterochromatin. Mol Cell. 2004;16:93–105.

29. Vaquero A, Scher MB, Dong HL, Sutton A, Cheng HL, Alt FW, et al. SirT2 is a histone deacetylase with preference for histone H4 Lys 16 during mitosis. Genes Dev. 2006;20:1256–1261.

30. Penney J, Tsai LH. Histone deacetylases in memory and cognition. Sci Signal. 2014;7:re12–re12.

31. Herskovits AZ, Guarente L. Sirtuin deacetylases in neurodegenerative diseases of aging. Cell Res. 2013;23:746–758.

32. Carafa V, Rotili D, Forgione M, Cuomo F, Serretiello E, Hailu GS, et al. Sirtuin functions and modulation: from chemistry to the clinic. Clin Epigenetics. 2016;8.

33. Bosch-Presegué L, Vaquero A. Sirtuin-dependent epigenetic regulation in the maintenance of genome integrity. FEBS J. 2015;282:1745–1767.

34. Chopra V, Quinti L, Kim J, Vollor L, Narayanan KL, Edgerly C, et al. The Sirtuin 2 Inhibitor AK-7 Is Neuroprotective in Huntington’s Disease Mouse Models. Cell Rep. 2012;2:1492–1497.

35. Biella G, Fusco F, Nardo E, Bernocchi O, Colombo A, Lichtenthaler SF, et al. Sirtuin 2 Inhibition Improves Cognitive Performance and Acts on Amyloid-β Protein Precursor Processing in Two Alzheimer’s Disease Mouse Models. J Alzheimers Dis. 2016;53:1193–1207.

36. Jing E, Gesta S, Kahn CR. SIRT2 Regulates Adipocyte Differentiation through FoxO1 Acetylation/Deacetylation. Cell Metab. 2007;6:105–114.

37. Wang F, Nguyen M, Xiao-Feng Qin F, Tong Q. SIRT2 deacetylates FOXO3a in response to oxidative stress and caloric restriction. Aging Cell. 2007;6:505–514.

38. Gearing LJ, Cumming HE, Chapman R, Finkel AM, Woodhouse IB, Luu K, et al. CiiiDER: A tool for predicting and analysing transcription factor binding sites. PLoS One. 2019;14:e0215495.

39. Nagashima T, Shigematsu N, Maruki R, Urano Y, Tanaka H, Shimaya A, et al. Discovery of novel forkhead box O1 inhibitors for treating type 2 diabetes: improvement of fasting glycemia in diabetic db/db mice. Mol Pharmacol. 2010;78:961–970.

40. Pandithage R, Lilischkis R, Harting K, Wolf A, Jedamzik B, Lüscher-Firzlaff J, et al. The regulation of SIRT2 function by cyclin-dependent kinases affects cell motility. J Cell Biol. 2008;180:915–929.

41. Pereira JM, Chevalier C, Chaze T, Gianetto Q, Impens F, Matondo M, et al. Infection Reveals a Modification of SIRT2 Critical for Chromatin Association. Cell Rep. 2018;23:1124–1137.

42. Gatta V, D’Aurora M, Granzotto A, Stuppia L, Sensi SL. Early and sustained altered expression of aging-related genes in young 3xTg-AD mice. Cell Death Dis. 2014;5:e1054.

43. Kimura KI, Aburai N, Yoshida M, Ohnishi M. Sanguinarine as a potent and specific inhibitor of protein phosphatase 2C in vitro and induces apoptosis via phosphorylation of p38 in HL60 cells. Biosci Biotechnol Biochem. 2010;74:548–552.

44. Allen M, Carrasquillo MM, Funk C, Heavner BD, Zou F, Younkin CS, et al. Human whole genome genotype and transcriptome data for Alzheimer’s and other neurodegenerative diseases. Sci Data. 2016;3:160089.

45. Mostafavi S, Gaiteri C, Sullivan SE, White CC, Tasaki S, Xu J, et al. A molecular network of the aging human brain provides insights into the pathology and cognitive decline of Alzheimer’s disease. Nat Neurosci. 2018;21:811–819.

46. Matarin M, Salih DA, Yasvoina M, Cummings DM, Guelfi S, Liu W, et al. A Genome-wide Gene-Expression Analysis and Database in Transgenic Mice during Development of Amyloid or Tau Pathology. Cell Rep. 2015;10:633–644.

47. Mucke L, Masliah E, Yu GQ, Mallory M, Rockenstein EM, Tatsuno G, et al. High-level neuronal expression of abeta 1-42 in wild-type human amyloid protein precursor transgenic mice: synaptotoxicity without plaque formation. J Neurosci. 2000;20:4050–4058.

48. Jankowsky JL, Fadale DJ, Anderson J, Xu GM, Gonzales V, Jenkins NA, et al. Mutant presenilins specifically elevate the levels of the 42 residue beta-amyloid peptide in vivo: evidence for augmentation of a 42-specific gamma secretase. Hum Mol Genet. 2004;13:159–170.

49. Ferrari ME, Bernis ME, McLeod F, Podpolny M, Coullery RP, Casadei IM, et al. Wnt7b signalling through Frizzled-7 receptor promotes dendrite development by coactivating CaMKII and JNK. J Cell Sci. 2018;131.

50. Chacón MA, Varela-Nallar L, Inestrosa NC. Frizzled-1 is involved in the neuroprotective effect of Wnt3a against Abeta oligomers. J Cell Physiol. 2008;217:215–227.

51. Zhang H, Hao Y, Yang J, Zhou Y, Li J, Yin S, et al. Genome-wide functional screening of miR-23b as a pleiotropic modulator suppressing cancer metastasis. Nat Commun. 2011;2:1–11.

52. Salpea P, Russanova VR, Hirai TH, Sourlingas TG, Sekeri-Pataryas KE, Romero R, et al. Postnatal development-and age-related changes in DNA-methylation patterns in the human genome. Nucleic Acids Res. 2012;40:6477–6494.

53. Li G, Luna C, Qiu J, Epstein DL, Gonzalez P. Role of miR-204 in the regulation of apoptosis, endoplasmic reticulum stress response, and inflammation in human trabecular meshwork cells. Investig Ophthalmol Vis Sci. 2011;52:2999–3007.

54. Wu F, Jiao J, Liu F, Yang Y, Zhang S, Fang Z, et al. Hypermethylation of Frizzled1 is associated with Wnt/β-catenin signaling inactivation in mesenchymal stem cells of patients with steroid-associated osteonecrosis. Exp Mol Med. 2019;51:1–9.

55. Bonnefont J, Tiberi L, Van Den Ameele J, Potier D, Gaber ZB, Lin X, et al. Cortical Neurogenesis Requires Bcl6-Mediated Transcriptional Repression of Multiple Self-Renewal-Promoting Extrinsic Pathways Article Cortical Neurogenesis Requires Bcl6-Mediated Transcriptional Repression of Multiple Self-Renewal-Promoting Extrinsic Pathways. Neuron. 2019;103:1096-1108.e4.

56. Cacabelos R, Carril JC, Cacabelos N, Kazantsev AG, Vostrov A V., Corzo L, et al. Sirtuins in alzheimer’s disease: SIRT2-related genophenotypes and implications for pharmacoepigenetics. Int J Mol Sci. 2019;20.

57. Wang Y, Yang J, Hong T-T, Sun Y, Huang H, Chen F, et al. RTN4B-mediated suppression of Sirtuin 2 activity ameliorates β-amyloid pathology and cognitive impairment in Alzheimer’s disease mouse model. Aging Cell. 2020;19.

58. Silva DF, Esteves AR, Oliveira CR, Cardoso SM. Mitochondrial Metabolism Power SIRT2-Dependent Deficient Traffic Causing Alzheimer’s-Disease Related Pathology. Mol Neurobiol. 2017;54:4021–4040.

59. Esteves AR, Palma AM, Gomes R, Santos D, Silva DF, Cardoso SM. Acetylation as a major determinant to microtubule-dependent autophagy: Relevance to Alzheimer’s and Parkinson disease pathology. Biochim Biophys Acta - Mol Basis Dis. 2019;1865:2008–2023.

60. Langlet F, Haeusler RA, Lindén D, Ericson E, Norris T, Johansson A, et al. Selective Inhibition of FOXO1 Activator/Repressor Balance Modulates Hepatic Glucose Handling. Cell. 2017;171:824-835.e18.

61. Hock C, Heese K, Hulette C, Rosenberg C, Otten U. Region-Specific Neurotrophin Imbalances in Alzheimer Disease. Arch Neurol. 2000;57:846.

62. Sampaio TB, Savall AS, Gutierrez MEZ, Pinton S. Neurotrophic factors in Alzheimer’s and Parkinson’s diseases: implications for pathogenesis and therapy. Neural Regen Res. 2017;12:549–557.

