## Supplementary Figures for "Epigenetic repression of Wnt receptors in AD: a role for Sirtuin2-induced H4K16ac deacetylation of Frizzled1 and Frizzled7 promoters"

### Figure S1: Downregulation of *Fzd1* and *Fzd7* in AD correlates with reduced levels of H4K16ac and the concomitant increase in Sirt2 at their promoters

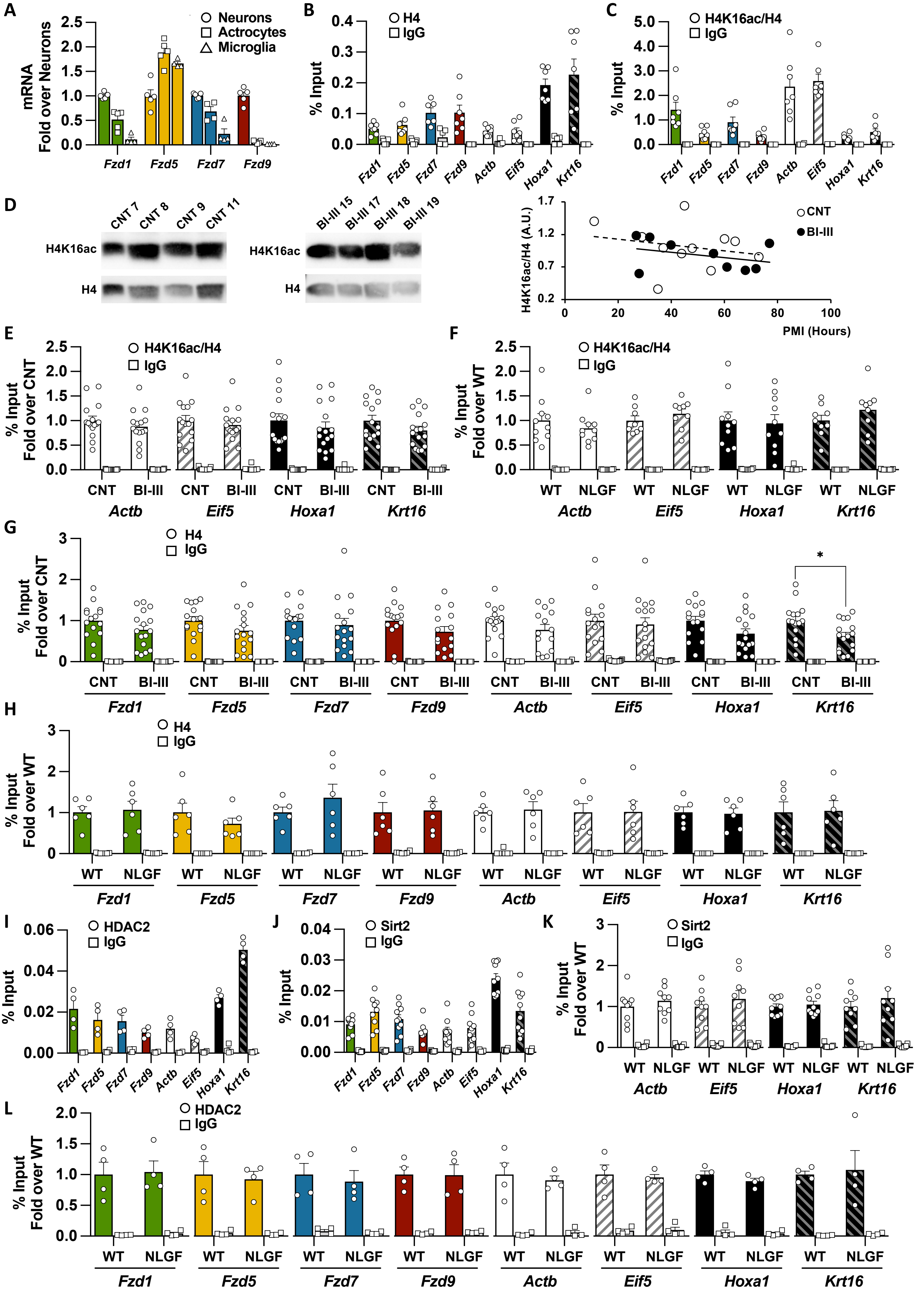

**Figure S1 legend: Downregulation of *Fzd1* and *Fzd7* in AD correlates with reduced levels of H4K16ac and the concomitant increase in Sirt2 at their promoters**

A) qPCR analyses of Fzds expression in neurons, astrocytes and microglia cultures. B) ChIP-qPCR analyses of total H4 in mouse hippocampal samples for the promoters of synaptic Fzds and the active (*Actb* and *Eif5*) and repressed (*Hoxa1* and *Krt16*) external control gens. Active control genes present low levels of H4, while repressed control genes present high levels of H4, correlating with their chromatin compaction and transcription in the brain. *Fzds* present intermediary levels of H4 at their promoters. C) ChIP-qPCR analyses of H4K16ac in mouse hippocampal samples. Active control genes present high levels of the pro-transcription histone mark H4K16ac, while repressed control genes present low levels of H4K16ac. *Fzd1* and *Fzd7* present higher levels of H4K16ac than *Fzd5* or *Fzd9* promoters. D) Representative WB image and correlational analyses of H4K16ac levels and post-mortem interval (PMI) in human control (CNT) and Braak I-II samples (BI-III) showing that PMI does not affect total H4K16ac levels. CNT: empty circles and dotted line, R=-0.227, p=0.502; BI-III: filled circles and continuous line, R=-0.351, p=0.355. E-F) ChIP-qPCR analyses of H4K16ac at the prompters of the external control genes *ACTB, EIF5, HOXA1* and *KRT16* in human Control and BI-III (E) and in WT and NLGF (F) samples showing no changes in the acetylation levels at the promoters of these four control genes in AD. G-H) ChIP-qPCR analyses of total H4 levels in human Control and BI-III (G) and WT and NLGF (H) samples, showing nor differences at promoters analysed. Reduced levels of total H4 are observed at *HOXA1* promoter (G), which is not translated into changes of H4K16ac levels in AD. I-J) ChIP-qPCR analyses of HDAC2 (I) and Sirt2 (J) in WT hippocampal samples showing no enrichment of these two H4K16 deacetylases at Fzds promoters in basal conditions. K) ChIP-qPCR analyses of Sirt2 at external control gene promoters in WT and NLGF hippocampal samples showing no changes for all genes analysed in AD. L) ChIP-qPCR analyses of HDAC2 at the prompters of *Fzds* and external control genes showing no changes in HDAC2 levels at promoters of all analysed genes. Data are represented as mean + SEM. Statistical analyses by *t*-Test in E for *ACTB*, *EIF5* and *KRT16* and by Mann-Whitney for *HOXA1*; in F *t*-Test for all genes; in G *t*-Test for *FZD1, FZD5, FZD9, ACTB, EIF5, HOXA1* and *KRT16* and by Mann-Whitney for *FZD7*; in H *t*-Test for all genes; in K *t*-Test for *Actb, Eif5* and *Krt16* and by Mann-Whitney for *Hoxa1*; in L *t*-Test for all genes. N are indicated in each bar by the number of symbols. Asterisks indicate $*p<0.05$.

#### Figure S2: Sirt2 role in regulating *Fzd* expression in neuronal cultures

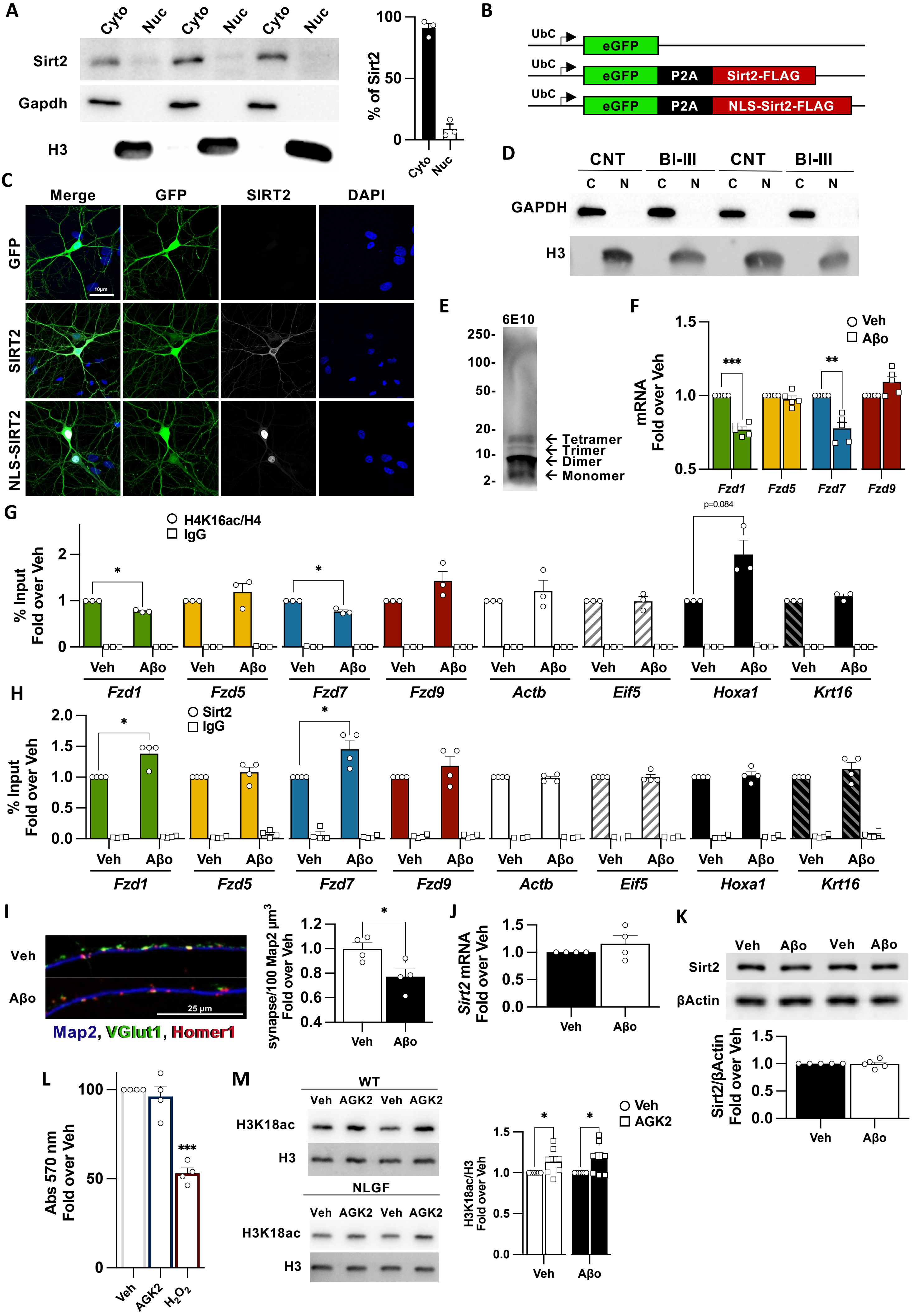

**Figure S2 legend: Sirt2 role in regulating *Fzd* expression in neuronal cultures**
A) Representative WB and quantification of Sirt2 in HEK cytosolic and nuclear fractions, showing that Sirt2 is mostly localised in the cytoplasm in HEK cells. B) Scheme representing the constructs used to overexpress WT Sirt2 or Sirt2 caring a nuclear localisation signal at its N-Terminal (NLS-Sirt2). All constructs contain GFP as a reporter. For Sirt2 constructs, GFP is followed by the cleaving peptide P2A, leading to the generation of GFP and Sirt2/NLS-Sirt2. C) Representative confocal images of the Sirt2 localisation; first column shows merged images with DAPI (blue), GFP (green) and Sirt2 (White), second column shows GFP, third column shows Sirt2 (white) and last column shows DAPI (blue). WT Sirt2 is mostly localised in the cytoplasm, while NLS-Sirt2 is mostly nuclear. D) Representative WB of nuclear enriched preparations for human CNT/BI-II showing clear fractionation of cytosolic and nuclear enriched extracts. E) Representative WB of the aggregative state of the synthetic Aβ oligomers (Aβo) used in our neuronal AD model. F) qPCR for *Fzd1, Fzd5, Fzd7* and *Fzd9* in neuronal cultures treated with 100 nM of Aβo O/N, showing reduced *Fzd1* and *Fzd7* expression. G-H) ChIP-qPCR experiments in neuronal cultures challenged with Aβo showing reduced H4K16ac (G) and a concomitant increase in Sirt2 (H) levels at *Fzd1* and *Fzd7* promoters. I) Representative image and quantification of synapses by the colocalization of the presynaptic marker vGlut1 (green) on the postsynaptic marker Homer1 (red) and Map2 (blue) in neuronal cultures challenged with Aβo. Our results show that Aβo leads to synapse loss. J-K) qPCR (J) and WB (K) for Sirt2 levels in vehicle (Veh) and Aβo treated cultures showing that Aβo does not change Sirt2 expression or protein levels. L) MTT cell viability analysis of the Sirt2 inhibitor AKG2 (7 μM) in neurons, showing no toxicity. As a positive control, neurons were challenged with 10 mM $H_2O_2$ for 2 hours. M) Representative WB and quantification of Veh and Aβo treated cultures with AGK2 showing increase acetylation of the Sirt2 substrate H3K18ac upon AGK2 treatment, suggesting effective Sirt2 inhibition. Data are represented as mean + SEM. Statistical analyses by one-sample *t*-Test in F for all genes; in G one-sample *t*-Test for *Fzd1*, *Fzd5*, *Fzd7*, *Fzd9*, *Actb*, *Eif5* and *Hoxa1* and by Mann-Whitney for *Krt16*; in H one-sample *t*-Test for *Fzd5*, *Fzd7*, *Fzd9*, *Actb*, *Hoxa1* and *Krt16* and by Mann-Whitney for *Fzd1* and *Eif5*;in I by *t*-Test; in J, K and L one-sample *t*-Test; in M one-tailed one-sample *t*-test for Veh and Aβo. N are indicated in each bar by the number of symbols. Asterisks indicate *$p<0.05$, **$p<0.01$, ***$p<0.005$.

### Figure S3: Sirt2 regulates *Fzd1* and *Fzd7* expression in organotypic cultures

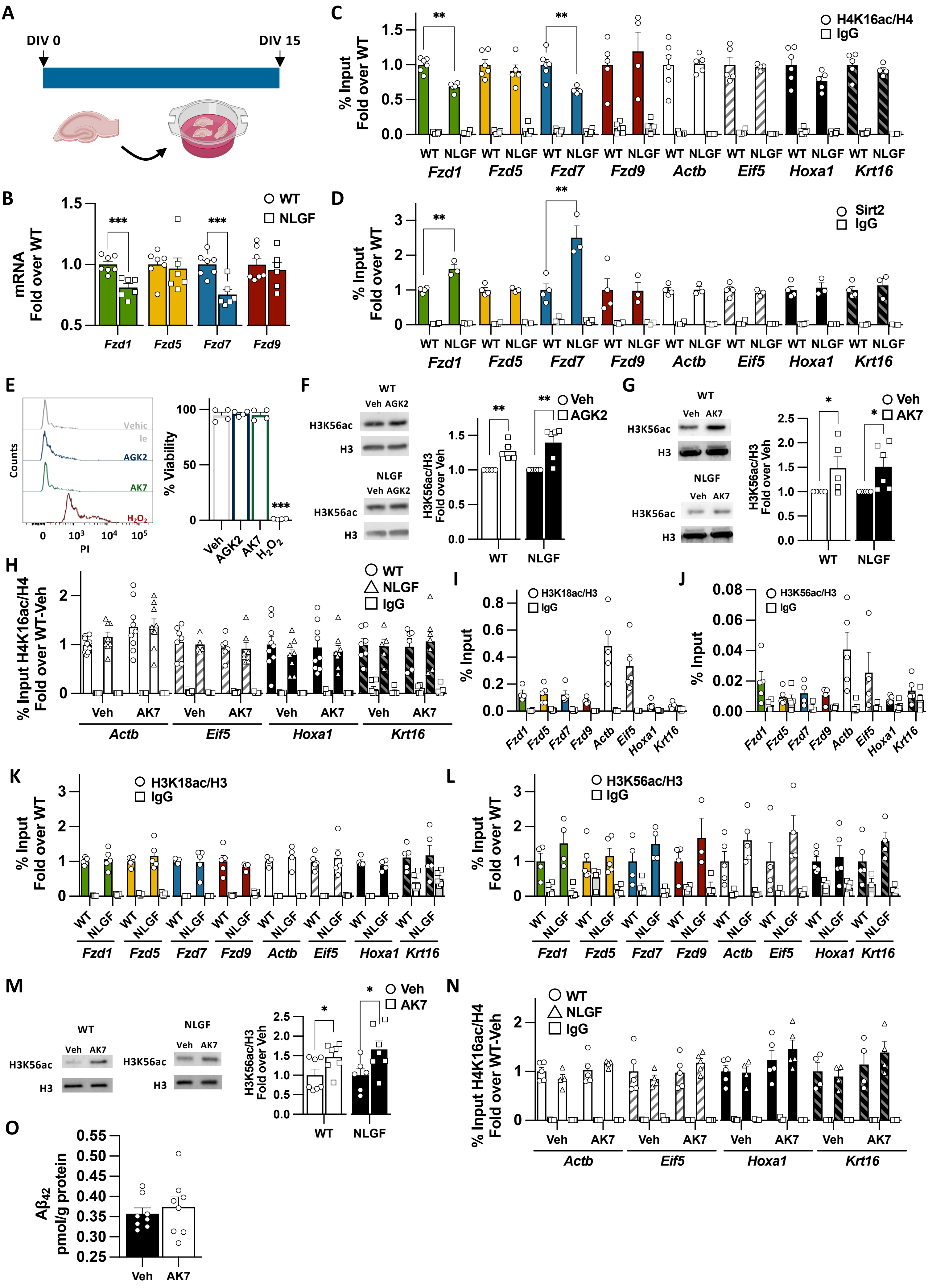

**Figure S3 legend: Sirt2 regulates *Fzd1* and *Fzd7* expression in organotypic cultures**

A) Scheme showing hippocampal organotypic cultures from WT and NLGF. B) qPCR results for 15DIV old hippocampal organotypic cultures showing the reduction in *Fzd1* and *Fzd7* levels in AD. *Fzd5* and *Fzd9* levels remain unchanged in AD after 15 days in culture. C-D) ChIP-qPCR experiments showing reduction in (C) H4K16ac and a concomitant increase in (D) Sirt2 at *Fzd1* and *Fzd7* promoters. H4K16ac and Sirt2 levels remain unchanged across the groups for the internal control gene *Fzd5* and *Fzd9* as for the external control genes *Actb, Eif5, Hoxa1* and *Krt16*. These results recapitulate the observations in human and NLGF at the RNA and epigenetic levels. E) Representative plots and quantification of cell viability FACS assay of WT hippocampal slices treated with vehicle, 7 µM AGK2, 30 µM AK7 or 10 mM $H_2O_2$ showing no cell toxicity for AGK2 or AK7. F-G) Representative WB and quantification of WT and NLGF treated with AGK2 (F) or AK7 (G) showing increase acetylation of the Sirt2 substrate H3K56ac upon AGK2/AK7 treatment, suggesting effective Sirt2 inhibition. H) ChIP-qPCR analyses showing that AK7 treatment does not affect the levels of H4K16ac at the external control genes *Actb, Efi5, Hoxa1* or *Krt16* promoters in WT and NLGF organotypic cultures. I-J) ChIP-qPCR analyses of H3K18ac (I) and H3K56ac (J) in WT hippocampal samples showing that these two Sirt2 substrates are not particularly enriched at *Fzd1* or *Fzd7* promoters. K-L) ChIP-qPCR experiments in WT and NLGF hippocampal samples for H3K18ac (K) and H3K56ac (L) showing no differential levels of these two histone marks at any of the promoters analyzed in AD. M) WB analyses of H3K56ac in frontal cortex samples of WT and NLGF animals treated with AK7 *in vivo* for 15 days. Our results show increased levels of H3K56ac thus suggesting the *in vivo* dosage of AK7 is effective in inhibiting Sirt2 in the brain. N) ChIP-qPCR analyses showing that AK7 treatment does not affect the levels of H4K16ac *in vivo* at the external control genes *Actb, Eif5, Hoxa1* or *Krt16* promoters in WT or AD. O) $A\beta_{42}$ quantification by ELISA in NLGF frontal cortex samples treated with vehicle (Veh) or AK7 showing no differences in $A\beta_{42}$ levels. Data are represented as mean + SEM. Statistical analyses by *t*-Test in B, C and D for all genes analysed; in E by *t*-Test to compare AGK2, AK7 or $H_2O_2$ to Vehicle; in F and G by one-tailed one-sample *t*-test for WT and NLGF; in H Two-way ANOVA followed by Tukey's post hoc for all genes analysed; in K and L by *t*-test for all genes analysed; in M one-tailed Mann-Whitney for WT and one-tailed *t*-test for NLGF; in N Two-way ANOVA followed by Tukey's post hoc for all genes analysed; in O *t*-test for $A\beta_{42}$ quantification. N are indicated in each bar by the number of symbols. Asterisks indicate *$p<0.05$; **$p<0.01$; ***$p<0.005$.

### Figure S4: Nuclear Sirt2 recruitment to *Fzd1* and *Fzd7* promoters in AD

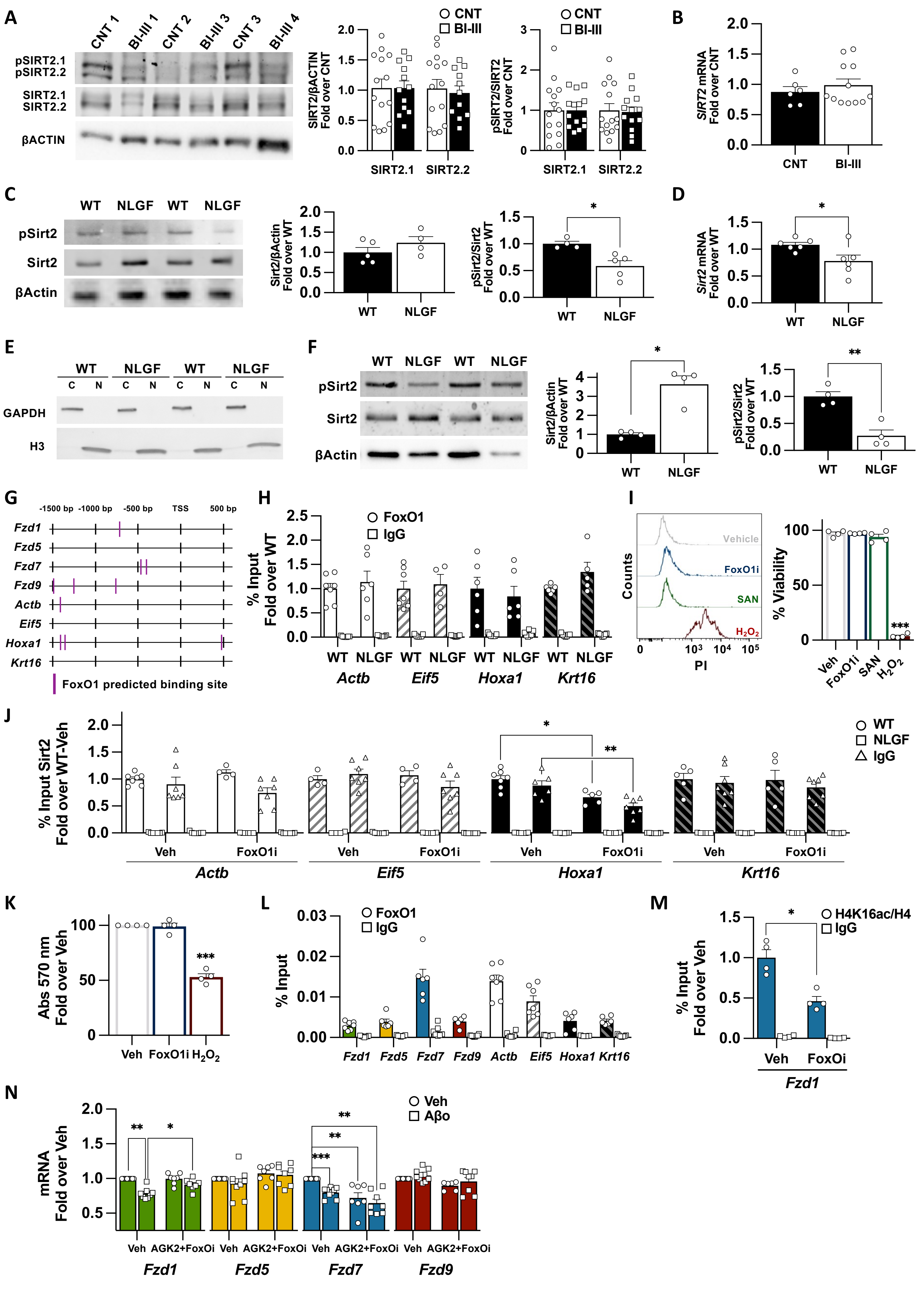

**Figure S4 legend: Nuclear Sirt2 recruitment to *Fzd1* and *Fzd7* promoters in AD**

A) WB analyses of total and pSIRT2 levels in hippocampal extracts of human control/BI-III subjects showing no differences of total or pSIRT2 at early AD. B) qPCR analysis of *SIRT2* expression in control/BI-III subjects showing no differences. C) WB analyses of total and pSirt2 levels in hippocampal organotypic cultures from WT and NLGF showing no differences of total Sirt2, but reduced pSirt2 in the AD model. D) qPCR analysis of *Sirt2* expression in WT/NLGF hippocampal organotypic cultures showing reduced *Sirt2* expression in NLGF. E) Representative WB of nuclear enriched preparations for organotypic WT/NLGF showing clear fractionation of cytosolic and nuclear enriched extracts. F) Representative WB and quantification for total and pSirt2 in nuclear extracts of WT/NLGF hippocampal organotypic cultures, showing increased Sirt2 levels and reduced pSirt2 levels in nuclear extracts of NLGF. G) Scheme of promoter region (-1500 bp –to TSS) and the start of the exons (TSS tot +500 bp) for *Fzd1*, *Fzd5*, *Fzd7, Fzd9*, *Actb, Eif5, Hoxa1* and *Krt16* containing CiiiDER predicted binding sites for FoxO1 in mouse. H) ChIP-qPCR analyses of FoxO1 at the prompters of the external control genes *Actb, Eif5, Hoxa1* and *Krt16* in WT and NLGF hippocampal samples, showing no changes in FoxO1 levels at the promoters of these four control genes in AD. I) Representative plots and quantification of cell viability FACS assay of WT hippocampal organotypic cultures treated with vehicle, 1 µM FoxO1I, 5 µM SAN or 10 mM $H_2O_2$ showing no cell toxicity for FoxO1I or SAN. J) ChIP-qPCR analyses showing that FoxO1i treatment does not affect the levels of Sirt2 at the promoters of the external control genes *Actb, Eif5 or Krt16* but reduces Sirt2 levels at *Hoxa1* promoters in both WT and NLGF organotypic cultures. K) MTT cell viability analysis for the FoxO1 inhibitor (1 µM) in neurons, showing no toxicity. As a positive control, neurons were challenged with 10 mM $H_2O_2$ for 2 hours. L) ChIP-qPCR analyses of FoxO1 in WT hippocampal samples showing FoxO1 is particularly enriched at Fzd7 promoter. M) ChIP-qPCR showing that FoxO1i treatment reduces H4K16ac levels at *Fzd7* promoters in WT organotypic cultures. N) qPCR analyses of Fzds expression upon co-inhibition of Sirt2 and FoxO1 by AGK2 and FoxO1i in vehicle (Veh) and Aβo treated neurons. Our results show that co-inhibition prevents *Fzd1* downregulation without modulating *Fzd5* or *Fzd9* mRNA levels. FoxO1 inhibition downregulates *Fzd7* expression *per se* and therefore, the co-inhibition elicits the same downregulation and fails to prevent its downregulation in Aβo treated neurons. Data are represented as mean + SEM. Statistical analyses in A by *t*-Test for pSIRT2.1 and by Mann-Whitney for total SIRT2.1, total SIRT2.2 and pSIRT2.2; in B by Mann-Whitney; in C *t*-Test, in F by Mann-Whitney for total Sirrt2 and by *t*-Test for pSirt2; in H by *t*-Test for all genes analysed, in I *t*-Test comparing FoxO1I, AK7 or $H_2O_2$ to Vehicle; in J Two-way ANOVA followed by Tukey's post hoc for *Eif5, Hoxa1* and *Krt16* and Kruskal-Wallis followed by Dunn's multiple comparison for *Actb*; in K by one-sample *t*-Test; in M by *t*-Test; in N Two-way ANOVA followed by Games-Howell post hoc for *Fzd1*, *Fzd5* and *Fzd7* and by Kruskal-Wallis followed by Dunn's multiple comparison for *Fzd9*. N are indicated in each bar by the number of symbols. Asterisks indicate *$p<0.05$; **$p<0.01$; ***$p<0.005$.

### Figure S5: Increased Sirt2 activity in AD impairs the transcription of synaptic Fzds

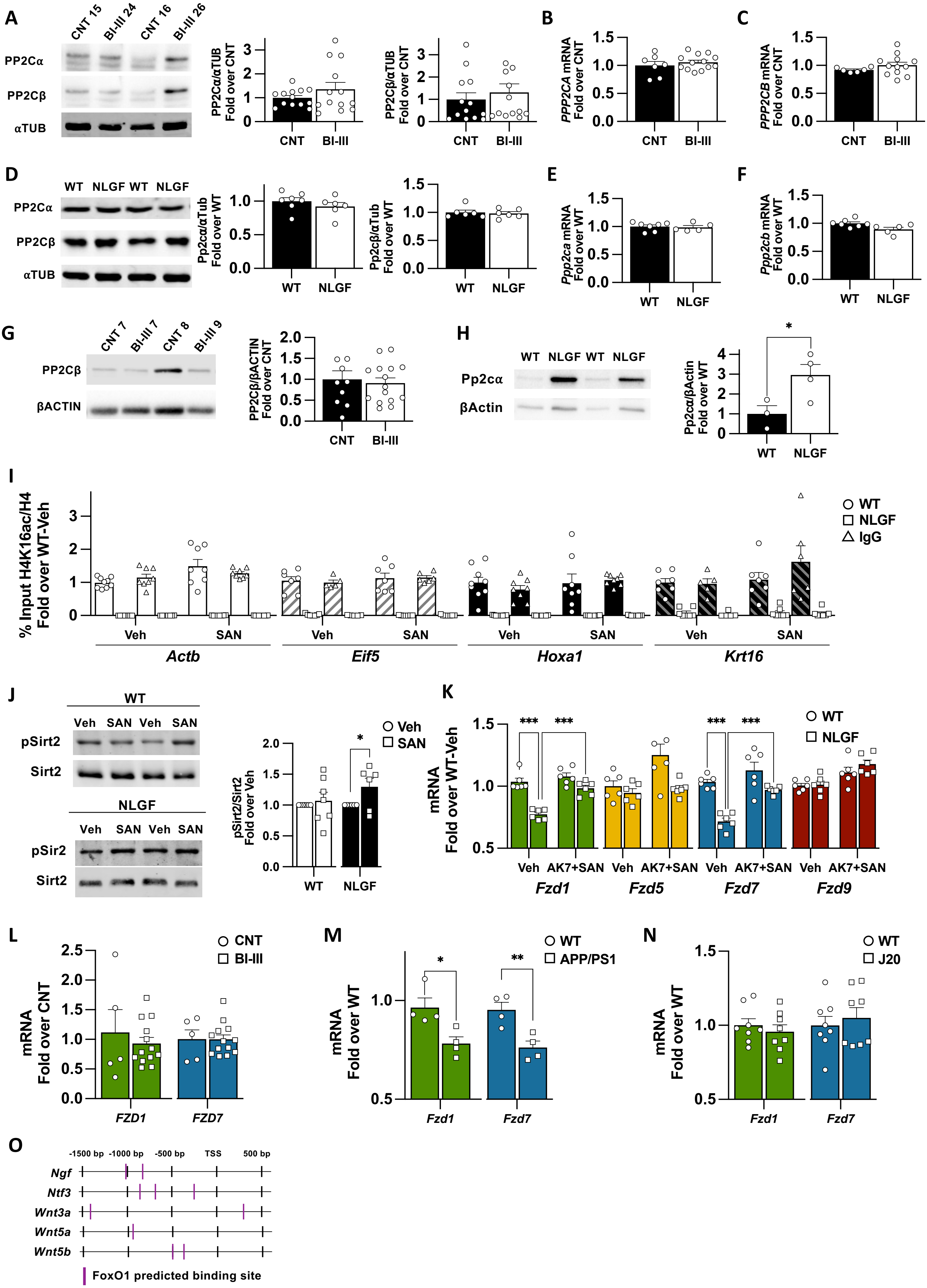

**Figure S5 legend: Increased Sirt2 activity in AD impairs the transcription of synaptic Fzds**
A) Representative WB and quantifications in total extracts from control/BI-III subjects showing no differences in total PP2Cα/β. B-C) qPCR analysis of *PPP2CA* (B) and *PPP2CB* (C) expression in control/BI-III subjects showing no differences. D) Representative WB and quantifications in total extracts from WT/NLGF hippocampal organotypic cultures showing no differences in total Pp2cα/β. E-F) qPCR analysis of *Ppp2ca* (E) and *Ppp2cb* (F) expression in WT/NLGF hippocampal organotypic cultures showing no differences. G) Representative WB and quantification of nuclear extract from control/BI-III subjects showing no differences in nuclear PP2Cβ. H) Representative WB and quantification for Pp2cα in nuclear extracts of WT/NLGF hippocampal organotypic cultures, showing increased nuclear Pp2cα levels in AD. I) ChIP-qPCR analyses showing that SAN treatment does not affect the levels of H4K16ac *in vitro* at the external control genes *Actb, Eif5, Hoxa1* and Krt16 promoters. J) Representative WB and quantification for pSirt2 in nuclear extracts of WT/NLGF hippocampal organotypic cultures treated with vehicle/SAN, showing increased nuclear pSirt2 levels only in NLGF. K) qPCR analyses of total mRNA levels from WT and NLGF hippocampal organotypic cultures treated with a co-inhibition of Sirt2 by AK7 and Pp2c by SAN. Our results show that AK7+SAN treatment rescues *Fzd1* and *Fzd7* mRNA levels and does not show any effect on *Fzd5* or *Fzd9* mRNA levels. L) qPCR analysis in prefrontal cortex samples from control/BI-III subjects showing no differences in *FZD1* or *FZD7* in this brain region. M-N) qPCR analysis in hippocampal samples from WT and the AD models APP/PS1 (M) or J20 (N), showing that *Fzd1* and *Fzd7* are downregulated at the hippocampus of APP/PS1 (M), but not in J20 (N). O) Scheme of the promoter region (-1500 bp –to TSS) and the start of the exons (TSS tot +500 bp) for *Ngf*, *Ntf3*, *Wnt3a*, *Wnt5a* and *Wnt5b* containing CiiiDER predicted binding sites for FoxO1 in mouse. Data are represented as mean + SEM. Statistical analyses in A by Mann-Whitney for Pp2cα/β; in B and C by *t*-Test; in D by *t*-Test for Pp2cα and Mann-Whitney for Pp2cβ; in E, F, G and H by *t*-Test; in I by Two-way ANOVA followed by Games-Howell post hoc for all genes analysed; in J by one-tailed one-sample *t*-test for WT and NLGF; in K by Two-way ANOVA followed by Tukey's post hoc for *Fzd1* and Fzd7 and by Two-way ANOVA followed by Games-Howell post hoc for *Fzd5 and Fzd7*; in L by *t*-Test for *FZD1* and *FZD7*; in M by *t*-Test for both genes; in N by *t*-Test for *Fzd1* and Mann-Whitney for *Fzd7*. N are indicated in each bar by the number of symbols. Asterisks indicate **p*<0.05; ***p*<0.01; ****p*<0.005.
