## Supplementary Methods for "Epigenetic repression of Wnt receptors in AD: a role for Sirtuin2-induced H4K16ac deacetylation of Frizzled1 and Frizzled7 promoters"

**SUPPLEMENTARY MATERIAL & METHODS**

**Human Tissue**

Anonymised human hippocampal samples from control and presymptomatic AD subjects were obtained from Cambridge Brain Bank (CBB), Division of the Human Research Tissue Bank, Addenbrooke’s Hospital, Cambridge UK. All samples were obtained with informed consent under CBB license (NRES 10/HO308/56). Anonymised human frontal cortex samples from control and presymptomatic AD subjects were obtained from Queen Square Brain Bank (QSBB) at UCL Queen Square Institute of Neurology. These samples were also obtained with informed consent under QSBB license (NRES 08/H0718/54). Tissues were stored at -80°C. Demographic data, Braak stages and experimental usage for each sample are shown in Table S1. Our hippocampal control cohort was composed by 50% male and 50% female with an average AAD of 62.12±4.43 SEM years. The hippocampal BI-III cohort was composed by 69.2% male and 30.8% female with an average age at death (AAD) of 78.88±2.23 SEM years. These demographic data could introduce a bias in our results as three control samples are younger (27, 35 and 37 years AAD) and the BI-III sample set was overrepresented with male samples. Our frontal cortex control cohort was composed by 60% male and 40% female with an average AAD of 80.0±3.39 SEM years. The frontal cortex BI-III cohort was composed by 30.8% male and 69.2% female with an average age at death (AAD) of 84.4±1.44 SEM years.

**Animals:**

All procedures involving animals were conducted according to the Animals Scientific Procedures Act UK (1986) and in compliance with the ethical standards at University College London (UCL).  Male and female 2 month-old hAPPNLGF/NLGF (NLGF) knock-in, 4 month-old APP/PS1 (B6-Cg-Tg(App_swe_, Psen1dE9)85Dbo-J), 4 month-old J20 (PDGF-APPSw,Ind) and corresponding WT littermates were used in this study. Genotyping was performed by PCR using DNA from ear biopsies. The following are the primers used for genotyping: For NLGF we used the following cocktail of primers: 5′-ATCTCGGAAGTGAAGATG-3′, 5′-ATCTCGGAAGTGAATCTA-3′, 5′-TGTAGATGAGAACTTAAC-3′ and 5′-CGTATAATGTATGCTATACGAAG-3′ [1]. For APP/PS1 we used three specific forward primers for Psen1 5′-CAGGTGGTGGAGCAAGATG-3’, App 5′-CCGAGATCTCTGAAGTGAAGATGGATG-3’ and PrP 5′-CCTCTTTGTGACTATGTGGACTGATGTCGG-3’, and one common reverse primer 5′-GTGGATACCCCCTCCCCCAGCCTAGACC)-3’ [2]. For J20 genotyping, we used 5’-GTGAGTTTGTAAGTGATGCC-3’ as forward and 5’-TCTTCTTCTTCCACCTCAGC-3’ as reverse [3]. Animals were kept in ventilated racks at 22±2°C and 55±10% humidity with a 12h/12h light cycle and free access to food and water at all times.

### *In vivo* AK7 treatment

For *in vivo* inhibition of Sirt2, WT and NLGF were treated with AK7. Given the limited brain availability of AK7 [4], animals were injected intraperitoneally twice a day (8 am and 18 pm) with 20 mg/Kg of AK7 (Tocris, ref. 4754), or the equivalent vehicle volumes, for 15 days. AK7 was diluted in DMSO (Sigma-Aldrich) to a final concentration of 40mg/mL, aliquoted and stored at -20 °C. AK7 aliquots, or equivalent DMSO volumes, were diluted to a final concentration of 4 mg/mL AK7, 10% DMSO and 25% Kolliphor® EL (Sigma-Aldrich) as previously described [4]. AK7 and vehicle were prepared fresh daily prior to injections. Animal wellbeing was monitored twice a day and body weights were recorded every three days. Samples were collected 3 hours after the last injection.

**Organotypic cultures**

Organotypic cultures were prepared from P9-P10 WT and NLGF animals following the protocol described by De Simony and Yu [5]. Briefly, hippocampi were extracted in slicing solution (EBSS (Thermo Fisher Scientific, ref. 24010-043) containing 25mM HEPES (Sigma-Aldrich, ref. H7523)) and sliced on an automatic tissue chopper (McIlwain Tissue Chopper, Standard Table, 220 V, Ted Pella Inc.) to obtain 400 μm hippocampal slices. Then, slices were separated in slicing solution. Finally, 5 hippocampal slices were placed onto Millicell culture inserts (Millipore, ref. PICM03050) in 6-weel plates containing 1 mL of equilibrated culture medium (50% MEM + Glutamax-1 (Thermo Fisher Scientific, ref. 42360-024), 18% EBSS (Thermo Fisher Scientific, ref. 24010-043), 25% Horse serum (Thermo Fisher Scientific, ref. 26050-070), 36 mM D-glucose (Sigma-Aldrich, ref G5767) and Pen/Step 1x (Thermo Fisher Scientific 15070-063)) and kept at 35 °C and 95% CO_2_ in a humidified incubator for 15 days. Culture medium was replaced in full every two/three days with equilibrated culture medium. Organotypic cultures were treated with 7 µM AGK2 (Sigma-Aldrich ref. 566324; stock 5mM in DMSO), 30 µM AK7 (Tocris ref.4754; stock 100 mM in DMSO), 100 nM AS1842856 (Millipore ref. 344355; stock 12.5 mM in DMSO) or 5 µM Sanguinarine (Tocris ref. 2302; stock 5 mM in DMSO). For drug specificities see Table S2 [6–8].

**Primary hippocampal cultures**

Primary cultures of hippocampal neurons were prepared from embryonic day 18 (E18) Sprague-Dawley rats as previously described [9]. Briefly, hippocampi were dissected and placed into ice-cold 1X Hank’s solution (LifeTechnologies ref.14185-045) with 7 mM HEPES (Sigma-Aldrich, ref. H7523). The tissue was then trypsinised (Life Technologies ref. 15090-046) at 37°C for 17 min. Hippocampi were washed three times with Hank’s solution. Cells were dissociated in 5 mL of plating medium (Dulbecco's Minimum Essential Medium (DMEM) high glucose, GlutaMAX supplemented with pyruvate (LifeTechnologies ref. 31966-021) supplemented with 10% horse serum (LifeTechnologies ref. 26050-088) and 20% glucose (LifeTechnologies ref. A24940-01)) and counted in a Neubauer Chamber. Cells were plated into pre-coated dishes with poly L-lysine (Sigma-Aldrich ref; P2636) for ChIP 750.000 cells in a 6 cm dish, for RNA 300.000 cells/well in a 6-well plate and for IF 170.000 cells/well in a 6-well plate containing 12 mm glass coverslips. Cells were placed into a humidified incubator containing 95% air and 5% CO_2_ at 37°C. The plating medium was replaced with equilibrated neurobasal media supplemented with B27, N2 and Glutamine (Gibco; Life Technologies refs.21103-049A35828-01, 17502-048). Neurons were cultured for 15 days in vitro (DIV). Cells where preincubated 30 min with relevant drugs (Table S2) prior to Aβ oligomers treatment O/N.

**Primary astrocyte and glial cultures**

Primary glial cultures were prepared from postnatal P1-P2 Sprague-Dawley rats as previously described [10]. Briefly, forebrain was dissected and placed into ice-cold 1X Hank’s solution (LifeTechnologies ref.14185-045) with 7 mM HEPES (Sigma-Aldrich, ref. H7523). The tissue was then chopped and trypsinised (Life Technologies ref. Life Technologies ref. 15090-046) and DNAse (Sigma-Aldrich E6267) and incubated at 37°C for 20 min. Tissue was washed three times with Hank’s solution. Tissue was dissociated in 10 mL of 10:10:1 medium (10% foetal bovine serum (LifeTechnologies ref.10108-165), 10% horse serum (LifeTechnologies ref. 26050-088) and 1% Pen/Strep (LifeTechnologies ref.15140-122)) and plated into a T75 flask pre-coated with poly L-lysine (Sigma-Aldrich ref. P2636). 24 hours later, medium was replaced with warm 10:10:1, mix glial cultures were grown for 7-10 days and then shacked at 300 rpm in an orbital shaker (Itertech ref 5003396) at 37°C. After 30 min shacking, medium containing microglia cells was recovered, and 10 mL of warmed 10:10:1 medium was added. Flasks were then shacked O/N. The following morning, medium was discarded and flasks were washed three times with warm PBS. Finally, astrocytes were detached with trypsin (LifeTechnologies ref.25200-056) for 5 min at 37°C. Microglia and astrocytes containing mediums were pellet for 5 min at 300 g, cells were resuspended in 1 mL of 10:10:1, counted in a Neubauer chamber and 600.000 cells were plated in a 6 cm dish pre-coated with poly L-lysine (Sigma-Aldrich ref. P2636). For astrocyte cultures, 2 hours after plating the 10:10:1 medium was replaced for IP-Astrocyte base medium (50% Neurobasal Medium (LifeTechnologies 21103), 50% DMEM (Life Technologies 11960-044), 1x Pen/Strep (LifeTechnologies 15140-122), 1 mM Sodium pyruvate (LifeTechnologies 11360-070), 5 µg/mL N-acetyl-L-cysteine (Sigma-Aldrich A8199), 5 ng/mL HBEGF (Sigma-Aldrich 4643) and SATO 1X (100 µg/mL BSA (Sigma-Aldrich A4161), 100 µg/mL Transferrin (Sigma-Aldrich T1147), 16 µg /mL Putrescine dihydrochloride (Sigma-Aldrich P5780), 0.2 µM Progesterone (Sigma-Aldrich P8783) and 40 ng/mL Sodium selenite (Sigma-Aldrich S5261))[11]. Cells were collected after 3-4 days in culture.

**Aβ aggregation**

Aβ oligomers (Aβo) were prepared as previously described [12] with minor modifications. Briefly, ultra-pure recombinant human Aβ_1-42_ HFIP film (Sigma-Aldrich ref. AG968) was resuspended at 500 µM in DMSO, sonicated in a Bioruptor-Pico (Diagenode, ref. B01060010) for 5 min (30 sedonds on/30 sedonds off) at room temperature, aliquoted and stored at -80°C. Aβ, or equivalent DMSO volume for vehicle, was diluted in neurobasal medium (Life Technologies ref. 21103-049) to a final 10 µM concentration, vortexed for 30 seconds and incubated at 4°C. 24 hours later, Aβo containing medium was vortexed, aliquoted and stored at -80°C. Aβo aliquots were thawed prior to use and the remainder was discarded. Each Aβ aggregation was evaluated by non-reducing SDS-PAGE: 40 µL of Aβo were loaded into a 4-20% Mini-PROTEAN TGX gel (Bio-Rad ref. 4561094), run, transferred onto a nitrocellulose membrane, blocked with 10% non-fat milk, incubated with the anti-Aβ antibody 6E10 (Table S4) and detected in a ChemiDoc MP Imaging System (Bio-Rad).

**Plasmids and transfection**

*Plasmids*: eGFP [13] and human *SIRT2*-FLAG [14] were introduced into FUW-UbC [15] backbone. Briefly, we amplified eGFP by PCR (Fw: 5’-GACAGGATCCGCCACCATGGTGAGCAAGGGC-3’, Rv: 5’-GCCCTCTAGATTACTTGTACAGCTCGTCCATGCC-3’), introducing BamHI restriction site at its 5’ end and stop codon followed by XabI restriction site at the 3’ end. This PCR product was digested and ligated into a FUW-ubiquitin backbone linearised with BamHI and XbaI to generate our control plasmid FUW-UbC-eGPF. Next, we amplified eGFP by PCR (Fw: 5’-GACAGGATCCGCCACCATGGTGAGCAAGGGC-3’, Rv: 5’-GGGCCCTCTAGAAGGCCCGGGGTTTTCTTCAACATCTCCTGCTTGCTTTAACAGAGAGAAGTTCGTGGCTCCGCTTCCCTTGTACAGCTCGTCCATGC-3’), introducing BamHI restriction site at its 5’ end and a P2A cleavage signal followed by XbaI restriction site at the 3’ end. This PCR product was digested and ligated into a FUW-UbC backbone linearised with BamHI and XbaI to generate and intermediary plasmid FUW-UbC-eGPF-P2A. WT SIRT2-FLAG was amplified by PCR (Fw: 5’-GGGCCTTCTAGAATGGACTTCCTGCGGAACTT-3’, Rv: 5’- GAATTCGGCGCGCCTTACTTGTCATCGTCGTCCTTGT-3’), introducing XbaI restriction site at its 5’ end and stop codon followed by AscI restriction site at the 3’ end. NLS-SIRT2-FLAG was amplified by PCR (Fw: 5’-CGGGCCTTCTAGAATGAAGCGGACTGCTGATGGCAGTGAATTTGAGTCCCCAAAGAAGAAGAGAAAGGTGGAAGGCGGCGACTTCCTGCGGAACTTATT-3’, Rv: 5’- GAATTCGGCGCGCCTTACTTGTCATCGTCGTCCTTGT-3’), introducing an XbaI restriction site and a BPSV40 nuclear localisation signal at its 5’ end and stop codon followed by AscI restriction site at the 3’ end. WT SIRT2-FLAG and NLS-SIRT2-FLAG PCR products were digested with XbaI and AscI and introduced into a FUW-ubiquitinn-eGPF-P2A backbone linearised with XabI and AscI to generate our experimental plasmids FUW-UbC-eGPF-P2A-SIRT2-FLAG and FUW-UbC-eGPF-P2A-NLS-SIRT2-FLAG.

*Transfection:* Plasmids were nucleofected using Amaxa Rat Neuron Nucleofector Kit (Lonza ref: VPG-1003). Briefly, 1.000.000 cells were pelleted for 5 min at 300 g and resuspended in 100 µL of Rat Neuron Nucleofector Solution. 1 µg of DNA was added, mixed, transferred into nucleofection cuvettes and nucelofected with program G-013. Nucleofecting solution canting cells was diluted by adding 900µL of plating medium. Finally, 400.000 cells/well were plated in 6-well plates pre-coated dishes with poly L-lysine (Sigma-Aldrich ref. P2636)

**RNA extraction**

Total RNAs from mouse and human hippocampal tissue or organotypic cultures was extracted with TRIzol Reagent (Thermo Fisher Scientific ref: 15596018) and DirectZol Miniprep RNA Kit (Zymo Research ref: R2052). Briefly, 20-50 mg of human hippocampal tissue or full mouse hippocampal tissue were homogenized in 800 µL of TRIzol using pellet pestles (Sigma-Aldrich ref: Z359947) coupled to pellet pestles cordless motor (Sigma-Aldrich ref: Z359971) in 1.5 mL tubes. 300.000 neurons or one insert for organotypic cultures were mechanically homogenised in 400µL of TRIzol by using a P1000 tip in 1.5 mL tubes. Once homogenized, samples were centrifuged for 5 min at 16,000g at 4 °C to remove any remaining tissue pieces and the supernatant was transferred to clean 1.5 mL tubes. 200 µL of chloroform for 1mL of TRIzol were added, tubes were vortexed for 5 sec and left at RT for 10 min. To obtain the phase separation, samples were centrifuged for 15 min at 10,000g at 4 °C. The aqueous upper face containing the RNA was transferred into a clean tube and RNA was extracted with DirectZol Miniprep RNA Kit following manufacturer’s instructions and including DNase in column treatment. RNA was quantified by measuring absorbance at 260 nm using a Nanodrop ND-100 (Thermo Fisher Scientific). RNA integrity (RNA Quality Indicator; RQI) was further analysed for human samples by using RNA HighSense Assay chips (Bio-Rad ref: 7007155) and run in Experion™ Automated Electrophoresis Station (Bio-Rad). Briefly, samples were diluted to 1-5 ng/µL, loaded into RNA HighSense Assay chips and run. 18S and 28S peaks were detected and RQI was determined. Only human samples with a RQI 3.9 or above were used for qPCR [15].

**Retrotranscription and qPCR**

Retrotranscription to first-strand cDNA was performed using RevertAid H Minus First Strand cDNA Synthesis kit (Themo Fisher Scientific ref: K1632). Briefly, up to 2000 ng of total RNA was used for cDNA synthesis following manufacturer’s instructions. 5 ng of original RNA was used to perform fast qPCR using GoTaq qPCR Master Mix (Promega ref: A6002) in a LigherCycler® 480 (Roche) following manufacturer’s protocol (2 min at 95°C followed by 40 cycles of denaturing (95°C) and annealing/extension (60°C)). Primers were designed using OligoPerfect design (Thermo Fisher Scientific) and *in silico* validated using *in silico* PCR (UCSC genome Browser) and Ensembl BLAST (Ensembl.org). Primers were purchased from Sigma-Aldrich (Table S3) and used at 0.5 μM final concentration. In previous work, we established the reference genes to use for mouse and culture samples (*Gapdh*, *Gusb* and *Pgk1*) [16], as well as for human samples (*PMU1*, *TBP*, *CYC1* and *UBE2D2*) [17].

**Single molecule Fluorescent in-situ hybridisation (smFISH)**

*Sample preparation*: Animals were induced by an overdose of Dolethal (Vetoquinol UK Limited, ref. 08007/4034) and subsequently perfused with cold PBS, followed by cold 4% PFA (Sigma-Aldrich, ref. P6148) diluted in PBS. Brains were extracted and placed in 4% PFA in PBS O/N, washed three times in cold PBS and placed in 30% sucrose in PBS O/N. Brains were frozen with isopentane (Sigma-Aldrich). 15 µm thick coronal brain sections were cut in a cryostat, placed onto a super-frost slide (Thermo Scientific) and stored at -80. Single molecule Fluorescent *in-situ* hybridisation (smFISH) was performed with RNAscope following manufacture instructions using RNAscope Multiplex Fluorescent V2 Kit (ACDBio, ref 323100). Briefly, slides containing frozen brains sections were dried in the oven at 40°C for 3-4min prior incubation in H_2_O_2_ for 10min at RT. Next, samples were boiled in antigen retrieval solution for 5 min, washed with 100% EtOH and dried for 5 min at RT. For antigen accessibility, slides were treated with Protease IV for 15 min at RT. Sections were washed twice in PBS. *Fzd7*-C2 (ACDBio, ref. 534101-C2) and Rbfxo3-C3 (ACDBio, ref. 313311-C3) probes were diluted in *Fzd1*-C1 probe (ACDBio, ref.404871) at a 1:50 ratio and incubated on the slides for 2hrs at 40°C. *Fzd1*-C1 probe was detected with TSA-Fluorescein 1:1000 (Perkin Elmer, ref. NEL741001KT), Fzd7-C2 probe with TSA-Cy3 1:1000 (Perkin Elmer, ref. NEL744001KT) and Rbfox3-C3 probe with TSA-Cy5 1:1000 (Perkin Elmer, ref. NEL745001KT). To quench the auto-fluorescence slides were incubated with TrueBlack (Biotium, ref.23007) for 30 sec at RT. Before mounting the slices, DAPI (Perkin Elmer, ref.323108) was added to stain nuclei. In this study, one positive (mouse *Ppib*, ACDBio, ref.320881) and one negative (Escherichia coli *DapB*, ACDBio, ref.320871) technical control probes were used. A one-day protocol was used in all experiments to preserve the quality of the slices.

*Image acquisition*: Images were acquired using a Zeiss 880 confocal microscope (x40 objective). Settings were established during the first acquisition and not modified afterwards. Stack images of 5 steps with a 0.5 µm interval were used. Two-three hippocampal sections per animal and 2-3 images per hippocampus were obtained at 1024x1024 pixels for image analysis. All images were pre-processed using ImageJ (maximum z-projection). *Image analysis*: Image analysis was performed using HALO (Indica Labs) and the image analysis pipeline described by Jolly et al. [17]. Briefly, colour deconvolution was performed, and excitation channels were assigned for detection of nuclear counterstains and mRNA probes (DAPI 405 nm, *Fzd1*-C1 probe 488 nm, *Fzd7*-C2 probe 568nm and Rbfox3-C3 647 nm). Nuclear segmentation and contour detection were optimised using the DAPI counterstain in real-time tuning mode using graphical overlays. Probe detection was optimised based on signal size (minimum signal size 0.5µm^2^), intensity of positive probe pixels and contrast threshold parameter settings. Probe signals were recorded both as number of individual fluorescent spots as well as total signal area in µm^2^ per cell on a continuous scale with conservative settings for spot segmentation and cluster counting (probe copy intensity threshold = 0.45, spot segmentation aggressiveness = 0.95). For absolute quantification of probe signals, physically separated puncta were always counted as one probe copy. A spot was classified as a cluster when its intensity surpassed the set probe copy intensity threshold. The probe count of each cluster is derived from absolute intensity divided by the copy intensity parameter. Thresholding for the copy intensity parameter was performed using individual probe signals, considering any background. Probe signals were assigned to each nucleus by proximity within a maximum cell radius of 25µm [18]. Total counts for the neuronal specific probe *Rbfox3* were used to determine cell identity (neuron/non-neuronal). We considered a positive neuronal cell when presenting 3 or more *Rbfox3* counts. Signal counts for the test probes *Fzd1* and *Fzd7* were quantified for each cell. The total number of probe counts associated with a cell were used to calculate % of *Rbfox3*^+^ and *Rbfox3*^-^ cells expressing 0, 1, 2, 3 or >3 copies of test probes *Fzd1* and *Fzd7* and H-scores. H-scores were calculated as follows: H-Score = (1×% Probe 1^+^Cells)+(2×% Probe 2^+^Cells )+(3×% Probe 3^+^Cells )+(4×% Probe >3^+^Cells), following recommendations by Cell Diagnostics USA (Newark, CA, USA) for the evaluation of RNAscope assays. Final scores derived by this metric has a range between 0 and 400.

**Chromatin immunoprecipitation (ChIP)**

ChIP experiments were performed as described by Palomer et al. [16]. Mouse tissue, 750.000 neurons or organotypic cultures were crosslinked with 1% formaldehyde for 15 min at room temperature. Crosslinking was stopped by adding glycine to a final concentration of 0.125 M for 2 min at room temperature. Samples were washed three times with cold PBS and homogenized in cold Soft Lysis Buffer (50 mM Tris (pH 8.0), 10 mM EDTA, 0.1% NP-40 and 10% glycerol) plus inhibitors (protease inhibitor (cOmplete, EDTA-free; Roche), phosphatase inhibitor cocktail 2 (Sigma-Aldrich) and the histone deacetylase inhibitor (NaBut at 5 mM; Sigma-Aldrich). Finally, lysates were centrifuged at 3,000 rpm at 4 °C for 15 min. Nuclei pellets were lysed with SDS Lysis Buffer (1% SDS, 10 mM EDTA and 50 mM Tris pH 8.0) plus inhibitors. Human tissues were homogenised with dounce homogenizer in 1 mL of STEN lysis buffer (1x STEN: 50 mM Tris, pH 7.6, 150 mM NaCl, 2 mM EDTA, 0.2% Nonidet P-40; STEN-lysis buffer, 1% Triton X-100, 1% Nonidet P-40, phosphatase and protease inhibitors in 1x STEN), and centrifuged at 4,000g for 5 minutes. Nuclei containing pellets were washed twice with STEN buffer and crosslinked with 1% formaldehyde in PBS, rotating for 15 min at RT. Crosslinking was stopped by adding glycine to a final concentration of 0.125 M, rotating for 2 min at room temperature. Samples were washed three times with cold PBS and homogenized in cold SDS Lysis Buffer containing phosphatase and protein inhibitors. Extracts were sonicated with Bioruptor-Pico (Diagenode, ref. B01060010) to generate DNA fragments below 1Kb. Samples were centrifuged at 13,000 rpm., 4 °C, 10 min to remove insoluble material, and the supernatant containing DNA–protein complexes was collected. The chromatin was diluted 1/10 with dilution buffer (0,01% SDS, 1,1% Triton X-100, 1,2 mM EDTA pH8, 16,7 mM Tris pH 8 and 167 mM NaCl) and pre-cleared with 50 μl protein A/G agarose beads (Santa Cruz Biotechnology, Inc.) and 20 μg of normal IgG (Santa Cruz Biotechnology, Inc.) for 500 μg of protein. Samples were placed in a wheel for 1–3 h at 4 °C. The mixture was centrifuged and the supernatant was collected. A total of 100 μg of protein or 1µg of DNA were used for each ChIP assay, reserving 10 % as the input. The antibodies (Table S4) were added to the chromatin lysate, incubated on a wheel O/N at 4 °C. Immune complexes were precipitated by the addition protein A/G agarose beads. As a negative control, non-immune IgG (Santa Cruz Biotechnology, Inc.) was used in place of specific antibodies. Immunoprecipitated complexes were washed three times with the following buffers: Low-salt wash buffer (0,1% SDS, 1% Triton X-100, 2 mM EDTA pH8, 20 mM Tris pH 8 and 150 mM NaCl); high-salt wash buffer (0,1% SDS, 1% Triton X-100, 2 mM EDTA pH8, 20 mM Tris pH 8 and 500 mM NaCl); and LiCl wash buffer (250 mM LiCl, 1% NP-40, 1% NaDOC, 1 mM EDTA and 10 mM Tris pH8). Immune complexes were eluted in 100 μl of 1% SDS and 100 mM NaHCO_3_ at 37 °C for 30 min. DNA–protein crosslinks were reversed by adding NaCl to a final concentration of 200 mM O/N at 60 °C. Protein digestion was performed 1 h at 37 °C by adding Proteinase K 0.04 mg/mL (Promega ref. MC5005), 50 mM EDTA pH8 and 500 mM Tris pH6,5 at final concentration. Finally, DNA was purified with QIAquick Gel Extraction Kit following the manufacturer procedures (Qiagen, Hilden, Germany) and eluted in 140 μl DNA/RNAse free MilliQ water. A total of 4 μl of purified DNA was used to perform fast qPCR using GoTaq qPCR Master Mix (Promega ref: A6002) in a LigherCycler® 480 (Roche) with the manufacturer’s protocol (2 min at 95°C followed by 40 cycles of denaturing at 95°C and annealing/extension at 60°C) and using the listed primers on Table S3. In all the cases, the ChIPs for histone marks have been normalized for total histone.

**Protein preparation and Western Blots**

Protein preparations from human, organotypic cultures and neuronal cultures were prepared following the same protocol. Briefly, total protein samples were mechanically lysed by using pellet pestles coupled to pellet pestles cordless motor in 1.5 mL tubes in RIPA buffer containing phosphatase and protein inhibitors and left 30 min on ice. Then samples were sonicated and insolubilized fractions were removed by centrifugation at 13,000g at 4 °C. Supernatants were collected and stored at -20 until use. Nuclear enriched fractions were prepared by homogenising samples with dounce homogenizer in 1 mL of STEN lysis buffer (1x STEN: 50 mM Tris, pH 7.6, 150 mM NaCl, 2 mM EDTA, 0.2% Nonidet P-40; STEN-lysis buffer, 1% Triton X-100, 1% Nonidet P-40, phosphatase and protease inhibitors in 1x STEN), and centrifuged at 4,000 rpm for 5 mi. Supernatants (Cytosol-enriched fraction) were recovered and stored at −20°C until use. Pellets (Nuclei-enriched fraction) were washed twice in ice-cold STEN buffer, solubilized in Tris 0.1M pH 8.0, SDS 1% containing protein and phosphatase inhibitors and sonicated. Protein quantification was determined with BCA Protein Assay (Thermo Scientific). Upon SDS-PAGE electrophoresis, proteins were transferred to nitrocellulose membranes and detected with the corresponding antibodies (Table S4) using a ChemiDoc MP Imaging System (Bio-Rad) or Odyssey CLx Imager (LiCor) developers and analysed by ImageJ.

**FACS cell viability**

Organotypic cultures were incubated with equilibrated culture medium containing 2 µg/mL of propidium iodide (PI) for 2 h (1mL under and 1 mL above the insert). Then, cultures were washed three times with PBS, lysed with STEN lysis buffer, lived for 30 min on ice and centrifuged at 4,000 rpm for 5 min. Nuclei enriched fractions were washed two times with PBS and incubated in PBS containing DAPI (Sigma Aldrich) for 5 minutes. Finally, samples were washed once with PBS, resuspended in FACS buffer (cold PBS plus 1% BSA) and passed through a 40µm cell strainer. All procedures/steps were performed at 4 °C. As positive control, cultures were treated with 10 mM of H_2_O_2_ for 15 min prior to PI incubation. Data were acquired with Diva acquisition software on a LSRII flow cytometer (Becton Dickinson) equipped with violet (405 nm), blue (488 nm), and red (633 nm) lasers. Post-acquisition analysis was performed using FlowJo software (TreeStar). PI levels were analysed on DAPI positive nuclei (gating on FSC/DAPI+). Threshold for PI positive dead cells was set up in H_2_O_2_ treated samples. Signal below threshold was quantified and expressed as percentage (%) of viability.

**MTT cell viability**

Neuronal cultures were grown for 15 DIV and treated with relevant drugs O/N (Table S2). Cell viability was measured by 3-[4,5-Dimethylthiazol-2-yl]-2,5-diphenyltetrazolium bromide (MTT) reduction (Sigma Aldrich ref. M5655). Briefly, 10 X MTT stock solution (5 mg/mL) was added to neuronal cultures 2 h prior to the end of the treatment. Next, medium was replaced with 1mL of DMSO and MTT reduction was determined in a plate reader spectrophotometer at 570 nm. Vehicle cells were taken as 100% viability. As a positive control, neurones were treated with 10 mM of H_2_O_2_ for 2h while incubated with MTT.

**Immunofluorescence: nuclear Sirt2 and Synapse quantification**

*Immunofluorescence staining:* Neuronal cultures were fixed in 4% PFA with 4% sucrose in PBS for 20 minutes, permeabilized with 0.05% Triton-X-100 in PBS for 5 minutes and blocked in 5% BSA for 1 hour, all at room temperature. Next, covers were incubated with primary antibodies (Table S4) O/N at 4°C. The following day, covers were washed three times with PBS and incubated with secondary antibodies (Table S4) for 1 hour at room temperature. Finally, covers were incubated with DAPI for 5 min (Table S4), washed three times with PBS and mounted with FluorSave (Millipore ref. 345789-20mL).

*Confocal microscopy:* 3 images per cover from three covers per culture were acquired on a Leica SP8 inverted confocal microscope. For synapse quantification: stacks comprised of 11 equidistant planes, 0.25 μm apart, were acquired using a 63x 1.40 NA oil objective. For Sirt2 overexpression and endogenous Sirt2 quantification: stacks comprised of 11 equidistant planes, 0.5 μm apart, were acquired using a 40x 1.30 NA oil objective.

*Image analyses:* For synapses, image analyses were performed using Volocity software (Perkin Elmer). Customized thresholding protocols were used to detect pre- and postsynaptic puncta (vGlu1 and Homer1) and Map2. Synapses were quantified as co-localized pre- and postsynaptic puncta on Map2. For nuclear Sirt2 quantification, image analyses were performed with Fiji. Briefly, GPF channel was used to generate a binary mask to quantify total Sirt2. The GFP mask was also used to select the corresponding cellular nucleus (DAPI staining) and subsequently generate a DAPI binary mask. In turn, the DAPI mask was used to quantify nuclear Sirt2. Nuclear Sirt2 was subtracted from total Sirt2 and expressed as % of cytosol/nuclear localization.

**Aβ_42_ Enzyme-linked immunosorbent assay (ELISA)**

Tissue samples were homogenised in RIPA buffer and soluble Aβ_42_ was quantified using the Human/Rat β-Amyloid_42_ ELISA Kit (Wako ref.  290-62601) following manufacturer’s instructions.

**Transcription binding sites analyses *in silico* by CiiiDER**

Putative binding sites for FoxO1 and FoxO3a at the promoter region (-1500 bp to TSS) and first 500 bp transcribed of different genes were analysed by the user-friendly transcription binding site predictor tool CiiiDER [19]. *In silico* analyses were performed in mouse genome (Mouse.GRCm38.94) using JASPAR2020_CORE_vertebrates.txt, which contains a curated, non-redundant set of profiles, derived from published and experimentally defined transcription factor binding sites for eukaryotes, with a deficit cut off threshold of 0.10.

**Supplementary References:**

1.	Saito T, Matsuba Y, Mihira N, Takano J, Nilsson P, Itohara S, et al. Single App knock-in mouse models of Alzheimer’s disease. Nat Neurosci. 2014;17:661–663.

2.	Knafo S, Sánchez-Puelles C, Palomer E, Delgado I, Draffin JE, Mingo J, et al. PTEN recruitment controls synaptic and cognitive function in Alzheimer’s models. Nat Neurosci. 2016;19:443–453.

3.	Tosh JL, Rickman M, Rhymes E, Norona FE, Clayton E, Mucke L, et al. The integration site of the APP transgene in the J20 mouse model of Alzheimer’s disease. Wellcome Open Res. 2017;2:84.

4.	Taylor DM, Balabadra U, Xiang Z, Woodman B, Meade S, Amore A, et al. A brain-permeable small molecule reduces neuronal cholesterol by inhibiting activity of sirtuin 2 deacetylase. ACS Chem. Biol., vol. 6, ACS Chem Biol; 2011. p. 540–546.

5.	De Simoni A, Yu LMY. Preparation of organotypic hippocampal slice cultures: Interface method. Nat Protoc. 2006;1:1439–1445.

6.	Carafa V, Rotili D, Forgione M, Cuomo F, Serretiello E, Hailu GS, et al. Sirtuin functions and modulation: from chemistry to the clinic. Clin Epigenetics. 2016;8.

7.	Nagashima T, Shigematsu N, Maruki R, Urano Y, Tanaka H, Shimaya A, et al. Discovery of novel forkhead box O1 inhibitors for treating type 2 diabetes: improvement of fasting glycemia in diabetic db/db mice. Mol Pharmacol. 2010;78:961–970.

8.	Kimura KI, Aburai N, Yoshida M, Ohnishi M. Sanguinarine as a potent and specific inhibitor of protein phosphatase 2C in vitro and induces apoptosis via phosphorylation of p38 in HL60 cells. Biosci Biotechnol Biochem. 2010;74:548–552.

9.	Dotti CG, Sullivan CA, Banker GA, Wong TP, Liu L, Lu J, et al. The establishment of polarity by hippocampal neurons in culture. J Neurosci. 1988;8:1454–1468.

10.	Miriam Mecha , Paula Marina Iñigo , Leyre Mestre , Miriam Hernangómez JB& CG. An easy and fast way to obtain a high number of glial cells from rat cerebral tissue: A beginners approach. : Protocol Exchange. Protoc Exch. 2011. https://www.nature.com/protocolexchange/protocols/2051. Accessed 11 June 2017.

11.	Foo LC. Purification of rat and mouse astrocytes by immunopanning. Cold Spring Harb Protoc. 2013;2013:421–432.

12.	McLeod F, Bossio A, Marzo A, Ciani L, Sibilla S, Hannan S, et al. Wnt Signaling Mediates LTP-Dependent Spine Plasticity and AMPAR Localization through Frizzled-7 Receptors. Cell Rep. 2018. 2018. https://doi.org/10.1016/j.celrep.2018.03.119.

13.	North BJ, Marshall BL, Borra MT, Denu JM, Verdin E. The human Sir2 ortholog, SIRT2, is an NAD+-dependent tubulin deacetylase. Mol Cell. 2003;11:437–444.

14.	Cortina C, Palomo-Ponce S, Iglesias M, Fernández-Masip JL, Vivancos A, Whissell G, et al. EphB-ephrin-B interactions suppress colorectal cancer progression by compartmentalizing tumor cells. Nat Genet. 2007;39:1376–1383.

15.	Rydbirk R, Folke J, Winge K, Aznar S, Pakkenberg B, Brudek T. Assessment of brain reference genes for RT-qPCR studies in neurodegenerative diseases. Sci Rep. 2016;6:37116.

16.	Palomer E, Carretero J, Benvegnù S, Dotti CG, Martin MG. Neuronal activity controls Bdnf expression via Polycomb de-repression and CREB/CBP/JMJD3 activation in mature neurons. Nat Commun. 2016;7.

17.	Jolly S, Lang V, Koelzer VH, Sala Frigerio C, Magno L, Salinas PC, et al. Single-Cell Quantification of mRNA Expression in The Human Brain. Sci RepoRtS |. 2019;9:12353.

18.	Fiala JC, Harris KM. Dendrite structure. 1999.

19.	Gearing LJ, Cumming HE, Chapman R, Finkel AM, Woodhouse IB, Luu K, et al. CiiiDER: A tool for predicting and analysing transcription factor binding sites. PLoS One. 2019;14:e0215495.
