## Supplementary Tables for "Epigenetic repression of Wnt receptors in AD: a role for Sirtuin2-induced H4K16ac deacetylation of Frizzled1 and Frizzled7 promoters"

**Table S1: Demographic data for human samples**

**A) Hippocampal Samples**

| Sample | Braak stage | Gender | Age at death | PMI (h) | RNA (RQI) | Experimental usage | | | |
|---|---|---|---|---|---|---|---|---|---|
| | | | | | | qPCR | ChIP | Total protein | Nuclear enriched fractions |
| CNT 1 | 0 | F | 85 | 29 | 3.5 | | X | X | X |
| CNT 2 | 0 | F | 54 | 37 | 3.6 | | X | X | X |
| CNT 3 | 0 | F | 72 | 38 | 3.2 | | X | X | |
| CNT 4 | 0 | M | 81 | 29 | 7.2 | X | X | X | |
| CNT 5 | 0 | F | 27 | 64 | 7.4 | X | X | X | X |
| CNT 6 | 0 | F | 60 | 60 | 6.9 | X | X | X | X |
| CNT 7 | 0 | M | 64 | 73 | 2.9 | | X | X | X |
| CNT 8 | 0 | F | 59 | 11 | 3.2 | | X | X | X |
| CNT 9 | 0 | M | 35 | 45 | 3.3 | | X | X | X |
| CNT 10 | 0 | M | 37 | 87 | 5.6 | X | X | X | |
| CNT 11 | 0 | M | 68 | 48 | 3.9 | X | X | X | X |
| CNT 12 | 0 | M | 73 | 31 | 3.8 | | X | X | |
| CNT 13 | 0 | F | 49 | 44 | 6.6 | X | X | X | X |
| CNT 14 | 0 | M | 83 | 45 | 5.0 | X | X | X | |
| CNT 15 | 0 | M | 78 | 55 | 3.4 | | X | X | X |
| CNT 16 | 0 | F | 69 | 35 | 3.8 | | X | X | X |
| BI-III 1 | I | F | 70 | 29 | 3.7 | | X | X | X |
| BI-III 2 | III | M | 87 | 44 | 3.2 | | | | X |
| BI-III 3 | II | M | 52 | 30 | 3.5 | | X | X | X |
| BI-III 4 | III | M | 79 | 46 | 6.1 | X | X | X | X |
| BI-III 5 | III | M | 90 | 17 | 6.0 | X | | | |
| BI-III 6 | II | M | 70 | 12 | 3.2 | | | | |
| BI-III 7 | II | F | 66 | 29 | 3.1 | | X | X | X |
| BI-III 8 | III | F | 77 | 55 | 5.8 | X | | X | |
| BI-III 9 | I | F | 86 | 40 | 3.5 | | X | X | X |
| BI-III 10 | III | F | 91 | 32 | 6.4 | X | X | X | X |
| BI-III 11 | II | F | 61 | 36 | 2.9 | | X | X | X |
| BI-III 12 | II | M | 83 | 27 | 5.7 | X | | X | |
| BI-III 13 | III | F | 85 | 91 | 3.4 | | X | | X |
| BI-III 14 | I | M | 66 | 44 | 6.1 | X | | | X |
| BI-III 15 | II | M | 80 | 40 | 5.4 | X | | | X |
| BI-III 16 | III | M | 88 | 28 | 3.5 | | | X | |
| BI-III 17 | II | F | 75 | 89 | 3.3 | | X | | X |
| BI-III 18 | II | M | 67 | 73 | 3.1 | | | X | X |
| BI-III 19 | III | M | 71 | 70 | 4.8 | X | X | | X |
| BI-III 20 | II | M | 72 | 55 | 6.2 | X | | X | |
| BI-III 21 | II | M | 78 | 49 | 5.5 | X | X | X | X |
| BI-III 22 | II | M | 87 | 61 | 5.7 | X | | | X |
| BI-III 23 | II | M | 96 | 56 | 6.6 | X | X | X | |
| BI-III 24 | III | M | 95 | 77 | 6.1 | X | | X | X |
| BI-III 25 | II | M | 84 | 68 | 2.9 | | | | X |
| BI-III 26 | II | M | 95 | 72 | 2.7 | | | X | X |

**B) Frontal Samples**

| Sample | Braak stage | Gender | Age at death | PMI (h) | RNA (RQI) |
|---|---|---|---|---|---|
| CNT 1 | 0 | M | 85 | 79 | 3.9 |
| CNT 2 | 0 | M | 77 | 40 | 7.3 |
| CNT 3 | 0 | F | 86 | 40 | 3.9 |
| CNT 4 | 0 | M | 84 | 79 | 7.7 |
| CNT 5 | 0 | F | 68 | 45 | 6.6 |
| BI-III 1 | 1 | F | 86 | 119 | 4.1 |
| BI-III 2 | 2 | F | 73 | 24 | 4.5 |
| BI-III 3 | 2 | F | 83 | 99 | 4 |
| BI-III 4 | 2 | M | 88 | 16 | 5.3 |
| BI-III 5 | 3 | F | 81 | 32 | 5.9 |
| BI-III 6 | 3 | M | 85 | 78 | 6.3 |
| BI-III 7 | 3 | F | 93 | 30 | 5 |
| BI-III 8 | 3 | F | 87 | 79 | 4.2 |
| BI-III 9 | 2 | M | 76 | 79 | 5.9 |
| BI-III 10 | 2 | F | 91 | 99 | 6.3 |
| BI-III 11 | 1 | F | 87 | 52 | 6.1 |
| BI-III 12 | 2 | F | 86 | 120 | 5.9 |
| BI-III 13 | 2 | M | 81 | 51 | 4.0 |

### Table S2: Drug specificity

| Drug | Abbreviation | Stock (Vehicle) | Treatment | Formula | $IC_{50}$ | Supp. References |
|---|---|---|---|---|---|---|
| AGK2 | AGK2 | 5 mM (DMSO) | 7 µM | $C_{23}H_{13}Cl_2N_3O_2$ | Sirt2 $IC_{50}$=3.5µM; Sirt1/3 $IC_{50}$>50µM | [6] |
| AK7 | AK7 | 100 mM (DMSO) | 30 µM | $C_{19}H_{21}BrN_2O_3S$ | Sirt2 $IC_{50}$=15.5µM; Sirt1/3 no inhibition at 50µM | [6] |
| AS1842856 | FoxO1i | 12.5 mM (DMSO) | 100 nM | $C_{18}H_{22}FN_3O_3$ | FoxO1 $IC_{50}$=0.33µM; Foxo3a/4 $IC_{50}$>1µM | [7] |
| Sanguinarine | SAN | 5 mM (DMSO) | 5 µM | $C_{20}H_{14}ClNO_4$ | PP2C $IC_{50}$=2.5µM; PP1 $IC_{50}$=42.5µM; PP2A $IC_{50}$>100µM; PP2B $IC_{50}$=77µM | [8] |

**Table S3: Primers list**

### Human

**qPCR**

| Gene | Forward | Reverse |
|---|---|---|
| ***FZD1*** | 5'- TCAAACACGTGCAAAAGAGC -3' | 5'- GGACCAGGCCAGTAAACTCA -3' |
| ***FZD5*** | 5'- TGGGGACTGTCTGCTCTTCT -3' | 5'- GCTGGGGAGAGACGGTTAG -3' |
| ***FZD7*** | 5'- CCCGTTGGTTGTTAATTTGG -3' | 5'- CCTCTGGCTTAACGGTGTGT -3' |
| ***FZD9*** | 5'- AAGATCGGGGTCTTCTCCAT -3' | 5'- AAGTCCATGTTGAGGCGTTC -3' |
| ***PPP2CA*** | 5'- TGCAATGAAGTAGTCGACACCT -3' | 5'- TGGCGATACTTTGGGTGCAT -3' |
| ***PPP2CB*** | 5'- AGCACTTGGAATCACGGGTT -3' | 5'- CAAGATGCGCATGACATGGG -3' |
| ***SIRT2*** | 5'- TACTTCATGCGCCTGCTGAA -3' | 5'- CCACCAAGTCCTCCTGTTCC -3' |
| ***PMU1*** | 5'- AGTGGGGGACTAGGCGTTAG -3' | 5'- GTTTTCATCACTGTCTGCATCC -3' |
| ***TBP*** | 5'- GCCCGAAACGCCGAATATAA -3' | 5'- AATCAGTGCCGTGGTTCGTG -3' |
| ***CYC1*** | 5'- CACGGAGGATGAAGCTAAGG -3' | 5'- GCATGAACATCTCCCCATCT -3' |
| ***UBE2D2*** | 5'- CAGCACAGTGTTCAGCAGGT -3' | 5'- TCATTTGGCCCCATTATTGT -3' |

**ChIP**

| Gene | Forward | Reverse |
|---|---|---|
| ***FZD1*** | 5'- AGCAAAAGCAACCAAGCCTG -3' | 5'- GAGGAAGTAGTGGTGGTGGC -3' |
| ***FZD5*** | 5'- ACAGTTCCAAGACAGCTCGG -3' | 5'- CCTGCAGCTAAGCGTCCTTA -3' |
| ***FZD7*** | 5'- GCAGTGTGACTGGGTTTCCA -3' | 5'- GCCCCAAGTTTCAGCCAAAC -3' |
| ***FZD9*** | 5'- CTGGGCGACAAGAGCAAAAC -3' | 5'- CAGACTGGGTCGCCTCATTT -3' |
| ***ACTB*** | 5'- GCCAAAACTCTCCCTCCTCC -3' | 5'- CTCTCCCCTCCTTTTGCGAA -3' |
| ***EIF5*** | 5'- GAGCCTCAGGAAGCAGAAGG -3' | 5'- CTCGTCATTGGTTCGCTTGC -3' |
| ***HOXA1*** | 5'- CTTCCCTTCTCACCTCTCGC -3' | 5'- CCTCCCCACCGTTCAATGAA |
| ***KRT16*** | 5'- ACCACCGAACTCGCATTTCT -3' | 5'- AGAGTTCCCACCCAGCTTTG -3' |

### Mouse

**qPCR**

| Gene | Forward | Reverse |
|---|---|---|
| ***Fzd1*** | 5'- CTCTTCACGGTGCTCACGTA -3' | 5'- GTAACAGCCGGACAGGAAAA -3' |
| ***Fzd5*** | 5'- ACATGGAACGATTCCGCTAC -3' | 5'- TCCCAGTGACACACACAGGT -3' |
| ***Fzd7*** | 5'- AGAGACAAAGCGGGAAACAA -3' | 5'- GGCTTTGCCTGTAAAAGCTG -3' |
| ***Fzd9*** | 5'- TCTGTCTCATGCAGGGAGTG -3' | 5'- CCTTCTGCCCCTTCTTATCC -3' |
| ***Ppp2ca*** | 5'- GCGTCCCTGACTTAGTCCAC -3' | 5'- CTTGCTTGCCAATTCACCCC -3' |
| ***Ppp2cb*** | 5'- CAAGGTCTCAGAGGAAGCCG -3' | 5'- GGGCAAGATCAGGCATTCCT -3' |
| ***Sirt2*** | 5'- CCACGGCACCTTCTACACAT -3' | 5'- TCACACCTGGGAGTTGCTTC -3' |
| ***Gapdh*** | 5'- CGTCCCGTAGACAAAATGGT -3' | 5'- TCAATGAAGGGGTCGTTGAT -3' |
| ***Gusb*** | 5'- GGTTTCGAGCAGCAATGGTA -3' | 5'- GCTGCTTCTTGGGTGATGTC -3' |
| ***Pgk1*** | 5'- TACCTGCTGGCTGGATGG -3' | 5'- CACAGCCTCGGCATATTTCT -3' |

**ChIP**

| Gene | Forward | Reverse |
|---|---|---|
| ***Fzd1*** | 5'- ACTTGTCCTCGCCCTTTCTT -3' | 5'- GGACAGGCTAGGTCGCTTTT -3' |
| ***Fzd5*** | 5'- GGTGGCACAACCGAATTGAC -3' | 5'- GTCCTTATCGAGGTCTGGCG -3' |
| ***Fzd7*** | 5'- CTCGAGAAACTTGGGCAAAC -3' | 5'- GACGAGCAACAAGCTTCTCC -3' |
| ***Fzd9*** | 5'- GGAAGAAAAGGGCGCACTTG -3' | 5'- GGAACGCACCTCCCAATGTA -3' |
| ***Actb*** | 5'- GAGACATTGAATGGGGCAGT -3' | 5'- ATGAAGAGTTTTGGCGATGG -3' |
| ***Eif5*** | 5'- GCGGTCCAGATGAGCAGTTA -3' | 5'- ATGTCCACGGTGCTTCTCTG -3' |
| ***Hoxa1*** | 5'- GCCACAAGAGAGCCAGGAG -3' | 5'- TGAACTGGCAAGAGGTGAGA -3' |
| ***Krt16*** | 5'- CTGGAGTCAGCAGTTGGAGG -3' | 5'- GTGCCTCAGACTCCAGGTTC -3' |

### Rat

**qPCR**

| Gene | Forward | Reverse |
|---|---|---|
| ***Fzd1*** | 5'- CCTGCGGACTGTAGAGGAAG -3' | 5'- GGGCAAAGCACTCATCAAAT -3' |
| ***Fzd5*** | 5'- TCTGTTATGTGGGCAACCAA -3' | 5'- CCAAGACAAAGCCTCGTAGC -3' |
| ***Fzd7*** | 5'- GCAGTGGCTGAAAAGACTCC -3' | 5'- CAGTTAGCATCGTCCTGCAA -3' |
| ***Fzd9*** | 5'- CAGCTCTCACTGGCTTTGTG -3' | 5'- AACTGGTGCCCAGTACCAAG -3' |
| ***Sirt2*** | 5'- GAAGGAGAAGGGGCTGCTG -3' | 5'- ATGTGTAGAAGGTGCCGTGG -3' |
| ***Gapdh*** | 5'- AGACAGCCGCATCTTCTTGT -3' | 5'- CTTGCCGTGGGTAGAGTCAT -3' |
| ***Actb*** | 5'- GGCTCCTAGCACCATGAAGA -3' | 5'- CTGGAAGGTGGACAGTGAGG -3' |

**ChIP**

| Gene | Forward | Reverse |
|---|---|---|
| ***Fzd1*** | 5'- CCCGCTTCAGAGCTAGCTAC -3' | 5'- GTGGTGGTGGCAGAAGCTAT -3' |
| ***Fzd5*** | 5'- CAAGCACCGGACCTCGATAA -3' | 5'- GTTCTGCTCTCTGAACGCCT -3' |
| ***Fzd7*** | 5'- AGCAAGAACGAGCCCTTTGA -3' | 5'- GCCTCCTCTGATTCTGGAGC -3' |
| ***Fzd9*** | 5'- GCCAGCCTGGGTTATCTGAG -3' | 5'- CTGCCCCTCTCCTCTCTTCT -3' |
| ***Actb*** | 5'- CATCGCCAAACTCTTCATCC -3' | 5'- GAGCGAGAGAGAAAGCGAGA -3' |
| ***Eif5*** | 5'- GCGGGGAACTGTGAGATTGA -3' | 5'- TGGTTATTGGCCACAGACCC -3' |
| ***Hoxa1*** | 5'- TCTTGCGCACTGTACATTCA -3' | 5'- CCTCCATAGGACCAGAGAAGAA -3' |
| ***Krt16*** | 5'- ACTCACTGCCTGAAGACACG -3' | 5'- CTGTTCCTTGAGCCTGAGGG -3' |

**Table S4: Antibodies list**

| Antibody | Raised in | Clonality | Conjuageted | ChIP | WB | IP | Origin & Ref |
|---|---|---|---|---|---|---|---|
| α-Tubulin | Rabbit | pAb | - | - | 1:1000 | - | Abcam ref. 4074 |
| β-Actin | Mouse | mAb | HRP | - | 1:5000 | - | Abcam ref. ab197277 |
| GAPDH | Rabbit | mAb | - | - | 1:25000 | - | Abcam ref. ab181602 |
| Histone H3 | Rabbit | pAb | - | 2 µg | - | - | Abcam ref. ab1791 |
| Histone H3 | Mouse | mAb | - | - | 1:1000 | - | Cell Signaling ref. 14269 |
| Acetyl-Histone H3 (Lys18) | Rabbit | pAb | - | 2 µg | - | - | Abcam ref. ab1191 |
| Acetyl-Histone H3 (Lys56) | Rabbit | pAb | - | 4 µg | - | - | Millipore ref. 07-677-I |
| Acetyl-Histone H3 (Lys56) | Rabbit | pAb | - | - | 1:1000 | - | Cell Signaling ref. 4243 |
| Histone H4 | Rabbit | pAb | - | 3 µg | - | - | Abcam ref. ab10158 |
| Histone H4 | Mouse | mAb | - | - | 1:1000 | - | Cell Signaling ref. 2935 |
| Acetyl-Histone H4 (Lys16) | Rabbit | pAb | - | 2 µg | 1:1000 | - | Millipore ref. 07-329 |
| Monomethyl-Histone H4 (Lys20) | Rabbit | mAb | - | 2 µg | - | - | Abcam ref. ab177188 |
| HDAC2 | Mouse | mAb | - | 10 µg | - | - | Abcam ref. ab12169 |
| PP2Cα | Mouse | mAb | - | - | 1:1000 | - | Abcam ref. ab14824 |
| PP2Cβ | Rabbit | pAb | - | - | 1:500 | - | Abcam ref. ab70804 |
| phospho-Sirt2 (Ser331) | Rat | mAb | - | - | 1:1000 | - | Active Motif ref. 61363 |
| Sirt2 | Rabbit | mAb | - | 5 µg | 1:1000 | 1:1000 | Abcam ref. ab211033 |
| Amyloid-β (6E10) | Mouse | mAb | - | - | 1:1000 | - | Biolegend ref. 803001 |
| vGlut1 | Guinea Pig | pAb | - | - | - | 1:5000 | Millipore ref. AB5905 |
| Homer | Rabbit | pAb | - | - | - | 1:1000 | Synaptic systems ref. 160003 |
| Map2 | Chicken | pAb | - | - | - | 1:1000 | Abcam ref. ab92434 |
| GFP | Chicken | pAb | - | - | - | 1:500 | Millipore ref. 06-896 |
| DAPI | - | - | - | - | - | 1:10000 | Sigma-Aldrich ref. D9542 |
| normal mouse IgG | Mouse | - | - | assay dependet | - | - | ThermoFisher ref. 10400C |
| normal rabbit IgG | Rabbit | - | - | assay dependet | - | - | ThermoFisher ref. 10500C |
| anti-Rabbit | Donkey | - | HRP | - | 1:5000 | - | GE Healthcare ref. NA934V |
| anti-Mouse | Sheep | - | HRP | - | 1:5000 | - | GE Healthcare ref. NA931V |
| anti-Rat | Goat | pAb | IRDye® 800RD | - | 1:10000 | - | Abcam ref. ab253031 |
| anti-Mouse | Donkey | pAb | Alexa 647 | - | 1:10000 | - | ThermoFisher ref. A-31571 |
| anti-Rabbit | Donkey | pAb | IRDye® 680RD | - | 1:10000 | - | Abcam ref. ab216779 |
| anti-Rabbit | Donkey | pAb | Alexa 568 | - | - | 1:600 | ThermoFisher ref. A10042 |
| anti-Guinea Pig | Goat | pAb | Alexa 488 | - | - | 1:600 | ThermoFisher ref. A11073 |
| anti-Rabbit | Donkey | pAb | Alexa 647 | - | - | 1:600 | ThermoFisher ref. A31573 |
| anti-Chicken | Goat | pAb | Alexa 568 | - | - | 1:600 | ThermoFisher ref. A11041 |
| anti-Chicken | Goat | pAb | Alexa 488 | - | - | 1:600 | ThermoFisher ref. A11039 |
